# Heightened empathic responses led by imbalanced cortico-amygdala circuit in *Shank3* autism mouse model

**DOI:** 10.1101/2025.11.14.688290

**Authors:** Sheng-Nan Qiao, Yong-Hui Jiang

## Abstract

Affective empathy is defined as individual percept and respond to another emotion appropriately, an essential building block for human society. Historically, aberrant empathy was considered as major characteristics of autistic individuals but recently it has been challenged. It required mechanistic understanding of the empathy process, especially how autistic individual percept other emotion. By evaluating the social fear transfer response, we reported a critical role of *Shank3* gene, one of the most replicated causative autism genes, in cortico-amygdala circuit in balancing and integrating affective empathy process using preclinical mouse models. We found that *Shank3* complete deletion mouse model exhibited unexpected exaggerated affective responses, which was encoded by defective neural dynamics of *Shank3*-deficient cingulate projecting amygdala circuit. This work for the first time showcased an interplay between gene and affective empathy responses in mouse and laid the groundwork for modeling empathy in autism and other neuropsychiatric disease.

## Main Text

Empathy—the capacity to perceive, understand, and respond to the emotional states of others—is a cornerstone of human social interaction and communication ^1,2^. Empathy encompasses both cognitive components, which involve understanding another’s perspective, and affective components, which reflect the automatic sharing or resonance of emotions encompassing multiple dimensions of social behavior ^3-6^. Atypical empathy has long been recognized as a hallmark of autism but remains incompletely understood ^7, 8-11,12^. Defining and quantifying empathy in autistic individuals remains difficult due to confounding by comorbid intellectual disability and communication impairments and the inherently subjective nature of questionnaire-based assessments ^13, 14^. Despite empathy’s fundamental role in human social behavior, the neural mechanisms underlying affective empathy—a spontaneous and evolutionarily conserved form of empathic response—remain poorly understood and represent a key target for understanding autistic and neurotypical individuals.

Emerging studies have recognized that many species, including laboratory rodents, can perceive the conspecific’s emotion^6^. Rodents exhibit forms of affective empathy, such as distress contagion and prosocial responses to conspecifics, offering a tractable system for mechanistic and circuit investigation^15-21^. Importantly, rodent models, especially genetically modified disease models like autism model offer unique advantages for mechanistic exploration, enabling the use of genetic manipulation, optogenetics, and *in vivo* imaging to map the circuitry and molecular substrates involved in empathic behavior ^22-24^. Adapted from human affective empathy studies, the behavioral paradigm with one subject observing another experiencing distress has been well established in mouse. It has been implicated that cortex regions including anterior cingulate cortex (ACC)^17,19,25-28^, insula^20,29,30^, and prefrontal cortex (PFC)^26,31,32^, thalamus^33^, hippocampus^21^, and nucleus accumbens (NAc) and basolateral amygdala (BLA) were involved in this affective processes^34,35^. However, it has not been studied whether and how dysfunction of those brain regions are involved in affective empathy process in autism mouse model.

Mutations in SHANK3, a synaptic scaffolding protein critical for excitatory synapse formation and function, are among the most penetrant genetic causes of autism spectrum disorder (ASD) and are representative of a larger class of neurodevelopmental disorders with autism due to synaptic dysfunction^36,37 38-40^. *Shank3*-deficient mice recapitulate deficits in social interaction ^41-53^, social memory^54^, social recognition^55^, and social preference^56,57^ and altered synaptic plasticity. However, while substantial research has focused on social interaction and cognition, the role of *Shank3* in empathy-related processes, such as emotional contagion and prosocial responding, remains largely unexplored. We have previously characterized *Shank3* exon 4-22 deletion mouse model (*Shank3*^Δ*e4-22*^), which completely eliminates SHANK3 protein and demonstrates robust face validity for ASD-related phenotypes ^58-60^. We hypothesize that *Shank3*^Δ*e4-22*^ mouse exhibit abnormal affective empathy and can serve as a powerful model to dissect the molecular and circuit mechanisms underlying social-affective processing.

### Loss of Shank3 causes increased social fear transfer and reduced social fear retrieval

To examine empathy related behaviors in the *Shank3*^Δ*e4-22*^ model of autism, we adapted a well-established 3-day fear transfer behavior paradigm ^27^. In this test, observation of a cage mate receiving repeated electrical shocks induces a freeze response indicative of fear in an observing mouse (**Fig. 1A**). We paired wild-type (WT), *Sh3^-/-^* and *Shank3*^Δ*e4-22*^ heterozygous (*Sh3^+/-^*) observers with WT cage mate demonstrators (**Fig. S1A-C**). *Sh3^-/-^* observers demonstrated enhanced observational fear as indicated by significantly increased incidence of freezes compared to WT and *Sh3^+/-^* observers during the fear transfer test (**Fig. 1B-C**). *Sh3^-/-^* observers demonstrated significantly reduced fear retrieval as determined by the lower incidence of freezes when returned to the testing chamber 24 hours later, compared to WT and *Sh3^+/-^*observers (**Fig. 1D-1E**). But *Sh3^+/-^* observers exhibited similar freeze behavior compared to WT during process, a similar pattern which was seen in our previous reports ^58-60^. As a result of this finding, all subsequent experiments were performed in *Sh3^-/-^* mice.

**Figure 1.**
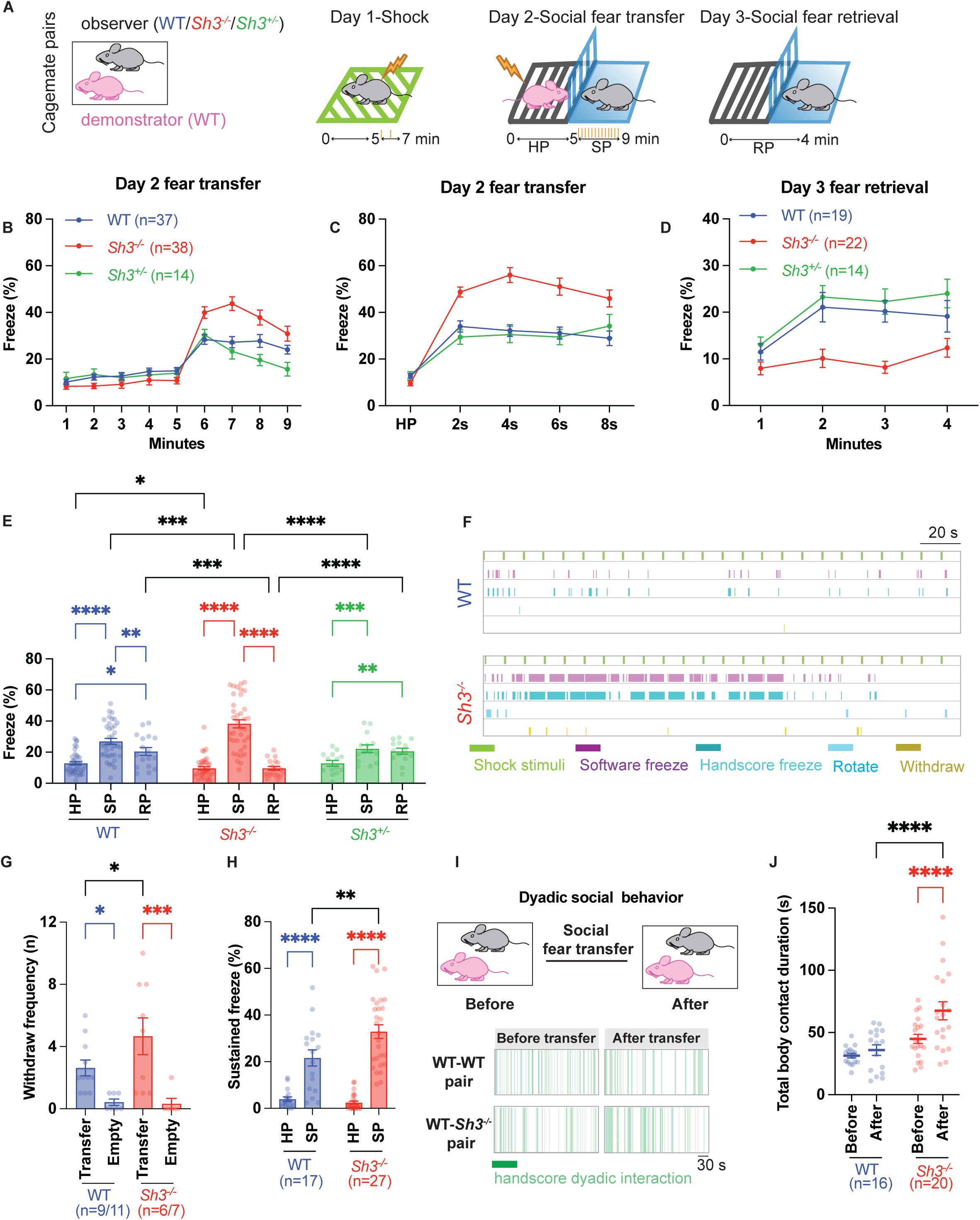
Sh3^-/-^ observer mouse exhibited heightened fear responses to social fear transfer compared to WT observer mouse. **A.** Schematic drawing showed the paradigm of social fear transfer behavior. Cage mate pairs of a WT demonstrator and a WT or *Sh3^-/-^* observer mouse was tested. At day 1, observer mouse received two mild electric shocks as prior experience exposure. At day 2, after 5-minute habituation period (HP), observer mouse observed demonstrator getting repetitive electric shocks during shocking period (SP) and observer’s freeze behavior was measured. At day 3, observer mouse returned to testing chamber with same context during retrieval period (RP) and observer’s freeze behavior was measured. **B-E.** Comparison of WT, *Sh3^-/-^* and *Sh3^+/-^* observer mouse’s freeze behavior among HP, SP, and RP phases. At day 2, WT, *Sh3^-/-^* and *Sh3^+/-^* observers significantly increased freeze responses in responses to demonstrator being shocked after 5-minute habituation. Compared to WT, *Sh3^-/-^*significantly increased freeze and *Sh3^+/-^* showed similar freeze, which was showed in both 1-minute (**B**) and 2-second (**C**) binned line plot. At day 3, compared toWT, *Sh3^-/-^* observer showed significantly reduced freeze while *Sh3^+/-^* observer showed similar freeze (**D**). On average, freeze duration of WT, *Sh3^-/-^* and *Sh3^+/-^*increased at SP than HP. Freeze duration of WT and *Sh3^+/-^*increased at RP than HP. Comparing WT vs. *Sh3^+/-^*, no significant difference of freeze duration was observed. But freeze duration of *Sh3^-/-^*significantly increased at SP and reduced at RP, compared to WT or *Sh3^+/-^*. *Sh3^-/-^* also showed decreased freeze duration than WT at HP. **F.** Examples of an individual WT and *Sh3^-/-^* observer mouse behavior at SP. Mice showed sporadically freeze in response to demonstrator being shocked. Mouse freeze was scored either automatically (software freeze) or manually (hand score freeze). Defensive behaviors such as rotation and withdraw were hand scored. **G.** Observer mice showed increased frequency of withdraw during fear transfer (transfer) compared to that in absence of demonstrator mouse (empty). *Sh3^-/-^* showed significantly increased frequency than WT. **H.** Observer mice increased sustained freeze responses (defined as one freeze episode longer than 2 seconds) at SP than HP. *Sh3^-/-^* showed significant longer sustained freeze duration than WT at SP. **I-J.** Example of dyadic free social interaction episodes from one pair of WT demonstrator and WT observer (WT-WT) and one pair of WT demonstrator and *Sh3^-/-^* observer (WT*-Sh3^-/-^*). Immediately before and after social fear transfer, social interaction was measured by hand scoring body contact duration (**I**). Pair of WT-*Sh3^-/-^* significantly increased social interaction duration after transfer than before, which is significantly longer than that from pair of WT-WT (**J**).

To further explore the observational fear phenotype of *Sh3^-/-^* observers, we hand scored freeze responses and analyzed innate defensive behavior primitives, including withdrawal, rotation, transient freeze, and sustained freeze ^61,62^ (**Fig. 1F, Fig. S1D, Fig. S2**). Withdrawal scores were significantly higher in the presence of a demonstrator being shocked (transfer) than in the absence of a demonstrator (empty), indicating that withdrawal truly reflected a fearful emotion. *Sh3^-/-^* observers exhibited more frequent withdrawal than WT, consistent with increased fear responses in *Sh3^-/-^* observers (**Fig. 1G**). Rotation frequency was similar between “transfer” and “empty” for both WT and *Sh3^-/-^* observers, although *Sh3^-/-^*observers displayed increased rotation frequency compared to WT (**Fig. S1E**). Overall, observers demonstrated more transient freezes than sustained freezes during the 5-min habituation period (HP) in the testing chamber before the onset of the shocking period (SP) although *Sh3^-/-^* observers exhibited fewer transient freezes during HP than WT. Overall, observers demonstrated more sustained freezes during SP, indicating that observers elongated freeze durations when experiencing fear transfer (**Fig. S1F**). *Sh3^-/-^* observers exhibited longer sustained freezes at SP compared to WT observers (**Fig. 1H**), indicating that *Sh3^-/-^* observers experienced increased fear (**Fig. S1F**). To elucidate if *Sh3^-/-^*observers adopted different social approaches and social communication during social fear transfer, we scored the average distance from observer to shocking section, duration observer spent sniffing the partition, and ultrasonic vocalization (USV) calls between the demonstrator-observer pair. WT and *Sh3^-/-^*observers showed similar social approach behaviors but WT-*Sh3^-/-^*pairs made fewer calls than WT-WT pairs (**Fig. S1G-I**). Because mice engage in prosocial behavior after stressful social contexts ^15,16,18^, we tested demonstrator-observer dyads for social interaction before and after fear transfer. WT-*Sh3^-/-^* pairs showed increased duration of social interaction after fear transfer, and also significantly longer social interaction after fear transfer compared to WT-WT pairs. This suggests that increased fear transfer in *Sh3^-/-^* observers facilitated subsequent social interactions in WT-*Sh3^-/-^* pairs (**Fig. 1I-J**).

Since it is reported female and male mice possess different sociability, we compared sex difference between genotypes. Female WT observers showed increased incidence of freeze responses compared to male WT observers, while female *Sh3^-/-^* observers showed decreased incidence of freeze responses compared to male *Sh3^-/-^*observers (**Fig. S1J-K**). The average freeze duration of male *Sh3^-/-^* observers was significantly longer than both WT males and WT female, while the average freeze duration of female *Sh3^-/-^* observers was significantly longer than WT males during the fear transfer test (**Fig. S1L**). Both WT and *Sh3^-/-^*Female observers showed significantly increased duration of social interaction after fear transfer, and increased duration compared to male observers, a general increased sociability often observed in female mice^63^. Additionally, Both male and female *Sh3^-/-^* observers showed significantly increased duration of interaction after fear transfer compared to WT observers in both sexes (**Fig. S1M**).

### Heightened social fear transfer due to loss of *Shank3* is independent of prior experience and depends on social familiarity

Previous studies reported that distinct neural circuits may underlie naive fear transfer (in absence of prior experience of shock) and experienced fear transfer (in presence of prior shock) in WT mice ^21,28^. Prior experience is modeled in the mouse fear transfer paradigm with a brief shock exposure to observer mice 1-day prior to the fear transfer test. To test if prior experience of shock is required for heightened fear transfer in *Sh3^-/-^* mice, we tested WT and *Sh3^-/-^* observers in a naive fear transfer paradigm (**Fig. 2A**). As expected, there was a reduced overall incidence of naïve observer freezes compared to experienced observers (**Fig. 1B-C**). However, naive *Sh3^-/-^*observers demonstrated significantly increased observational fear compared to naïve WT observers (**Fig. 2B-C**). WT and *Sh3^-/-^* observers demonstrated similar incidences of retrieval freeze responses when returned to same chamber 24 hours later (**Fig. 2D**). Freeze duration during SP was longer, and freeze duration during RP was shorter, in naive *Sh3^-/-^* observers than naïve WT observers (**Fig. 2E**). The *Sh3^-/-^* enhanced observational fear phenotype remained largely unchanged between the experienced and naive fear transfer tests, indicating that social fear hypersensitivity in *Sh3^-/-^*observers was sustained in the absence of early experience exposure.

**Figure 2.**
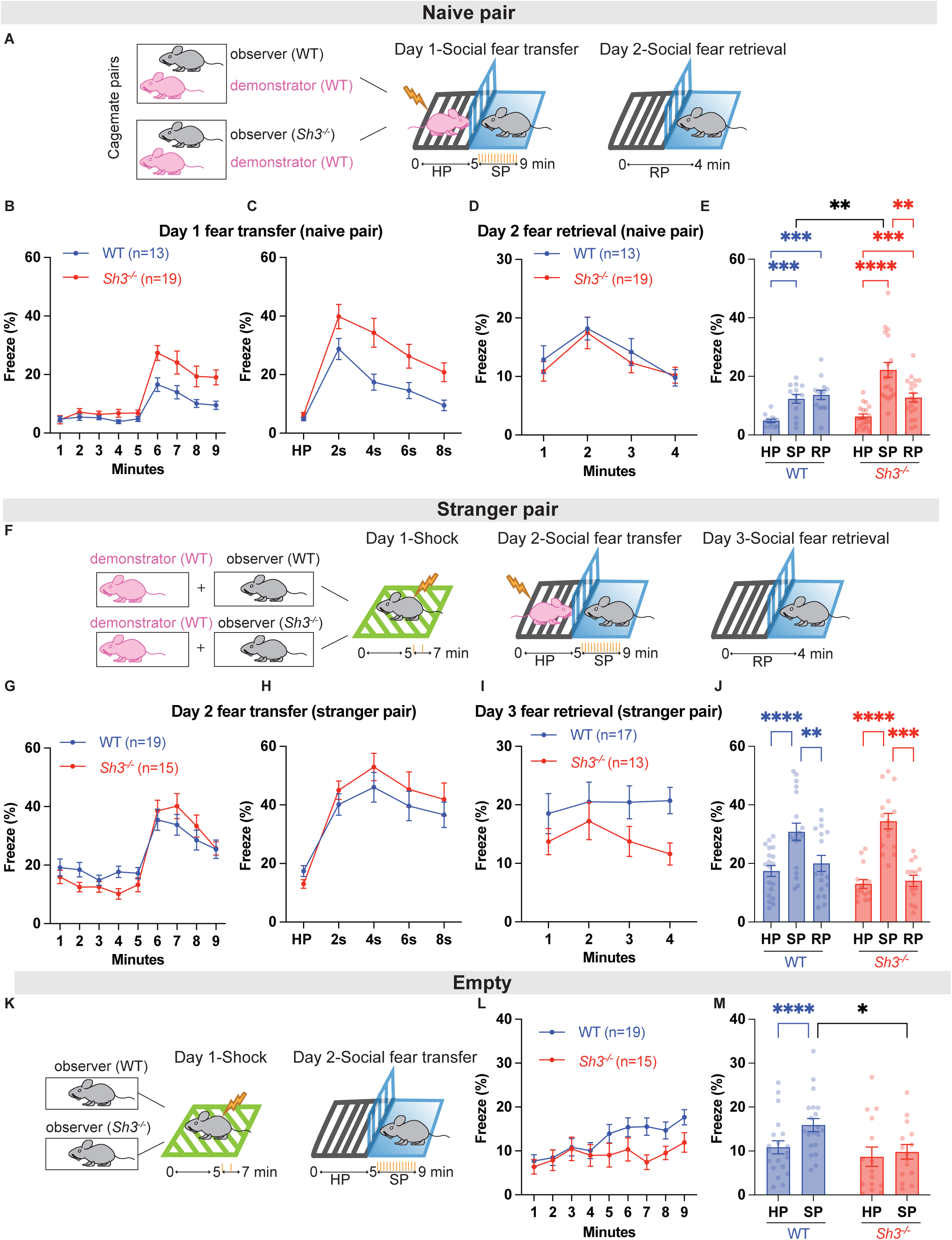
Sh3^-/-^ observer mouse heightened social fear transfer response was dependent on social familiarity and independent on prior experience. **A-E.** *Sh3^-/-^* observer mouse showed heightened social fear transfer response in absence of prior experience. Schematic drawing showed the behavior paradigm for naïve pair. Cage mate pairs of WT-WT and WT-*Sh3^-/-^* were tested. At day 1, observer mouse observed demonstrator getting shocked and freeze response was measured. At day 2, observer mouse returned same test chamber and freeze response was measured (**A**). Observer mice significantly increased freeze in responses to demonstrator being shocked after 5-minute habituation but *Sh3^-/-^* observers showed significantly increased freeze than WT at day 1 (**B-C**). *Sh3^-/-^* observers showed similar freeze responses as WT at day 2 (**D**). On average, both WT and *Sh3^-/-^* observers significantly increased freeze at SP and at RP than HP. *Sh3^-/-^* observers showed significantly increased freeze than WT at SP (**E**). **F-J.** Stranger pair of WT-WT and WT-*Sh3^-/-^* showed similar social fear transfer responses. Schematic drawing showed the behavior paradigm for stranger pair. Demonstrator mouse and observer mouse from different cages were used for 3-day test of social fear transfer behavior (**F**). Observer mice showed similar freeze responses at day 2, although *Sh3^-/-^*observers showed slightly decreased freeze at day 3 (**G-I**). On average, WT and *Sh3^-/-^* observers significantly increased freeze responses at SP than HP and showed similar freeze responses at SP (**J**). **K-M.** Schematic drawing showed the behavior paradigm for empty assay. At day 1, WT or *Sh3^-/-^*mouse received mild shocks then were placed in the social fear transfer chamber in absence of demonstrator at day 2 (**K**). WT mouse slightly increased freeze whereas immobility of *Sh3^-/-^*mouse remained same during 9-minute assay (**L**). On average, without demonstrator mouse, WT mouse increased freeze duration at 5-9 minutes (SP) than 0-5 minutes (HP) but *Sh3^-/-^* mouse showed similar freeze duration at SP and HP. Compared to WT, *Sh3^-/-^* mouse showed lower spontaneous immobility at SP (**M**).

As a complex social emotion, empathy is underpinned by social familiarity in humans ^64,65^. To elucidate if social familiarity modulates affective empathy in *Sh3^-/-^* mice, we conducted social fear transfer tests with stranger mice paired from different cages (**Fig. 2F**). Strikingly, *Sh3^-/-^*observers demonstrated loss of observational fear hypersensitivity in the stranger fear transfer test (**Fig. 2G-H**). *Sh3^-/-^* observers demonstrated modestly lower incidences of retrieval fear responses than WT (**Fig. 2I**). Average freeze duration was similar between WT and *Sh3^-/-^*observer responses at HP, SP and RP (**Fig. 2J**). This suggests that social fear hypersensitivity in *Sh3^-/-^* was dependent on the social relationship between observer and demonstrator.

To determine whether habituation-related immobility confounded freeze responses, we measured freeze behaviors in an empty control experiment in which demonstrator mice were absent during whole process (**Fig. 2K**). Over the course of the 9-minute test, freeze duration of WT observers slightly increased due to reduced movement in the chamber and *Sh3^-/-^* observers maintained consistent low mobility (**Fig. 2L**). Freeze duration during minute 5-9 (SP) increased in WT observers compared to minute 0-5 (HP), while freeze duration didn’t change in *Sh3^-/-^*observers (**Fig. 2M**). These results show that habituation-related immobility does not confound heightened social fear transfer responses in *Sh3^-/-^* observers.

### *Shank3* regulates region-specific bidirectional modulation of social fear transfer responses

Because ACC and BLA regions have been implicated in empathy processing in both human and mouse studies ^34,35^ and SHANK3 protein is reported to be expressed in those regions ^57^, we investigated the role of ACC and BLA regions in heightened social fear transfer in the *Sh3^-/-^*mouse. We used a conditional *Shank3* allele (*Shank3* ^e4-22 flox/flox^, *Sh3^f/f^*) together with viral-expressed Cre recombinase to specifically target *Shank3* exons 4-22 for deletion via focal virus injection into the ACC and BLA. Efficiency of Cre-lox recombination-induced *Shank3* deletion and *Shank3* expression was confirmed in brain tissue after behavioral testing (**Fig. S3**). Targeting *Shank3* in the ACC diminished observational fear (**Fig. 3A**) but did not change fear retrieval (**Fig. 3B-D**) compared to injection of control virus expressing EGFP (**Fig. 3E**). In contrast, targeting *Shank3* in the BLA (**Fig. 3F**), increased observational fear and fear retrieval compared to injection of control EGFP virus (**Fig. 3G-H**). These results indicate a dual bidirectional regulatory role for *Shank3* in the ACC and BLA, suggesting that altered social fear transfer responses in *Sh3^-/-^* mice are driven by disrupted equilibrium in the ACC-BLA circuit.

**Figure 3.**
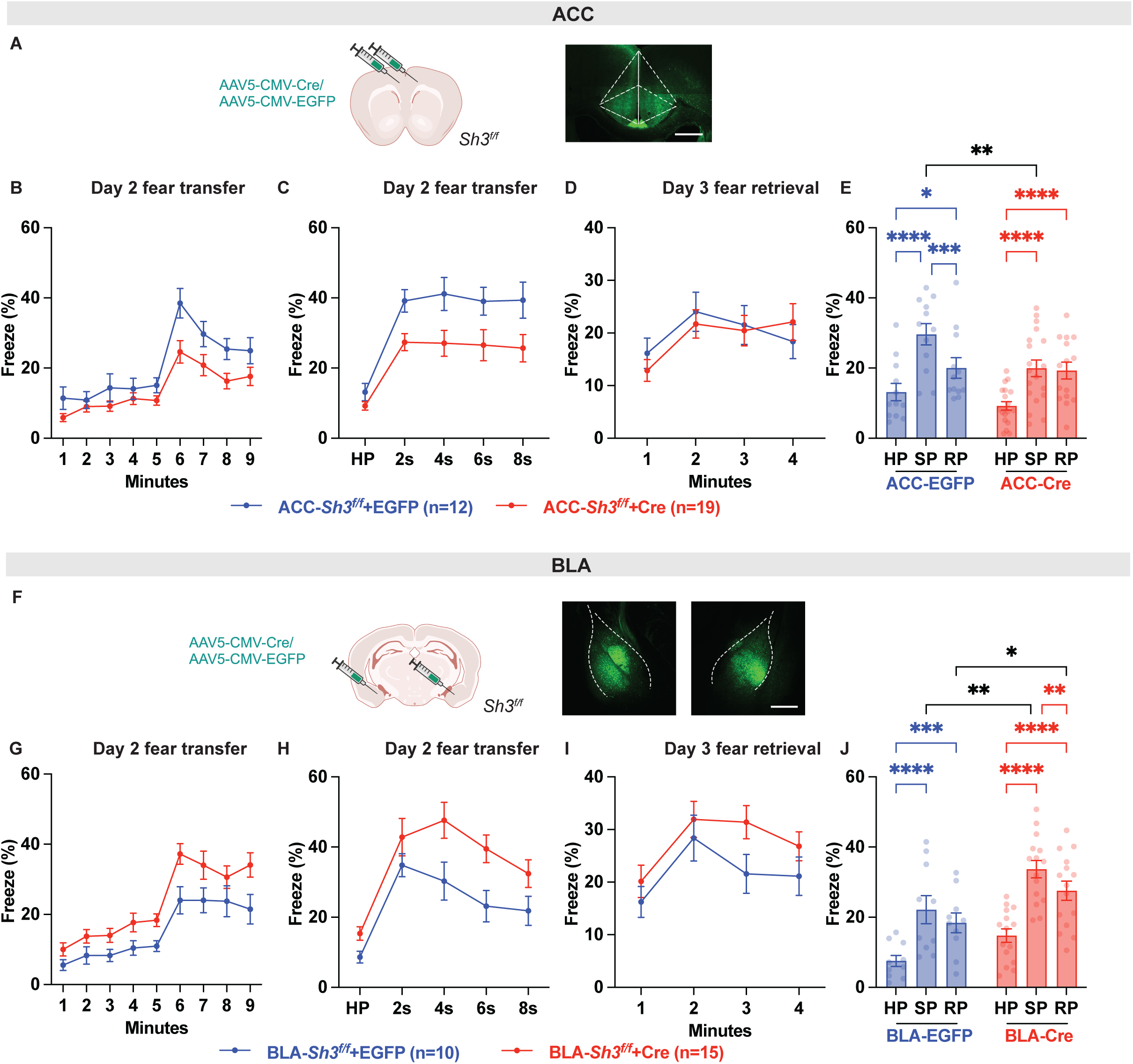
Shank3 controlled balance of social fear transfer process in ACC-BLA circuit. **A.** Schematic drawing showed knockdown of *Shank3* in *Sh3^f/f^* mouse via injecting AAV-Cre-EGFP or AAV-EGFP viruses in ACC. Example images showed virus expression in ACC. Scale bar is 500 μm. **B-E.** Conditional deletion of *Shank3* in ACC diminished social fear transfer. Compared to control group (ACC-*Sh3^f/f^*+EGFP), cre group (ACC-*Sh3^f/f^*+Cre) showed significant decreased freeze responses at SP but no change at RP. **F.** Schematic drawing showed knockdown of *Shank3* in *Sh3^f/f^* mouse via injecting AAV-Cre-EGFP or AAV-EGFP viruses in BLA. Example images showed virus expression in BLA. Scale bar is 500 μm. **G-J.** Conditional deletion of *Shank3* in BLA enhanced social fear transfer process. Compared to control group (BLA-*Sh3^f/f^*+EGFP), cre group (BLA-*Sh3^f/f^*+Cre) showed significant increased freeze responses at both SP and RP.

To confirm the efficiency of conditional manipulation of *Shank3* in brain region, we examined Cre-lox recombination induced *Shank3* deletion by extracting DNA and mRNA from virus-injected brain tissue. Primer pair KO was designed to detect deletion of *Shank3* gene between 1^st^ loxp and 2^nd^ loxp sites, and primer pairs ANK, PDZ and e21 were designed for reverse transcription quantitative PCR (RT-qPCR) to assess *Shank3* gene knockdown efficacy (**Fig. S3A**). DNA from Cre not EGFP virus-injected tissue successfully amplified using primer pair KO, suggesting Cre-induced deletion of *Shank3* gene (**Fig. S3B**). RT-qPCR showed that Cre virus induced ∼48% knockdown in ANK region, ∼21% knockdown in PDZ region and ∼25% knockdown in e21 region of *Shank3* gene (**Fig. S3C**). These results suggested successful conditional knock-down of SHANK3 in specific brain regions.

### Deficits of social fear-induced ACC and BLA neuronal activity and dynamics due to loss of Shank3

Our results that *Shank3* expression in the ACC is essential for normal social fear transfer and *Shank3* expression in the BLA is autonomous for heightened social fear transfer indicate an important function for *Shank3* in the ACC-BLA circuit in social fear transfer processes. To understand neuron activity dynamics during social fear transfer, we measured *in vivo* Ca^2+^ activity in ACC and BLA neuron populations in WT and *Sh3^-/-^* observers during social fear transfer. We injected a virus expressing GCaMP, a fluorescent calcium indicator, and simultaneously implanted a fiber optic cannula for fluorescence recording, in the ACC and BLA, respectively (**Fig. 4A-B**).

**Figure 4.**
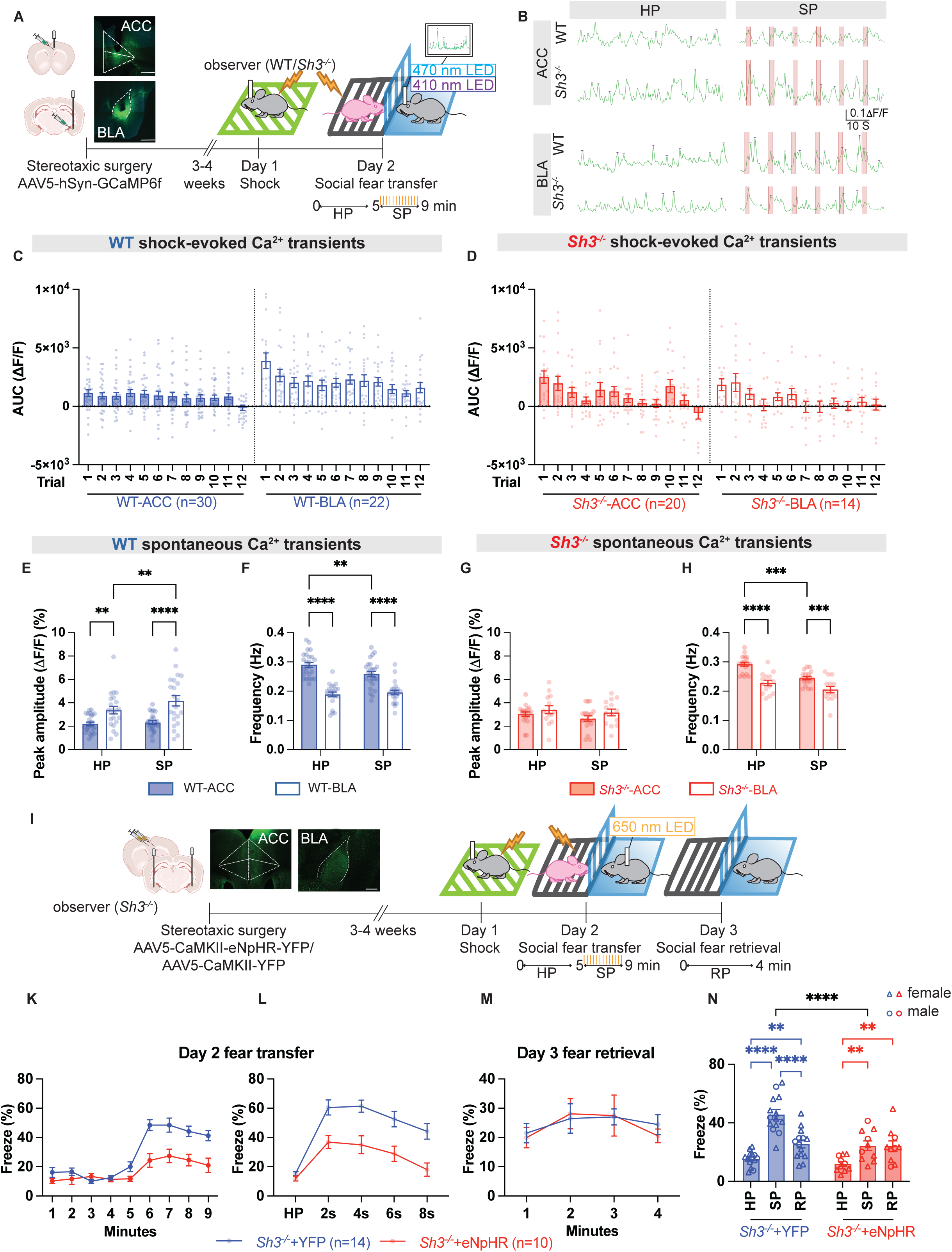
Sh3^-/-^ observer mouse heightened social fear transfer response was driven by defective neurons dynamics in ACC-BLA circuit. **A.** Schematic drawing showed *in vivo* Ca^2+^ imaging recording in ACC or BLA from observer mouse using fiber photometry recording. GCaMP virus expression and fiber optic cannula implant in ACC and BLA were validated post-mortem. Scale bar is 500 μm. **B.** Representative 1-minute window of Ca^2+^ signals in ACC and BLA from WT and *Sh3^-/-^*observer mouse at HP and SP. Shock stimuli delivered to demonstrator mouse was indicated as red bar. **C.** Area under curve (AUC) of shock event-aligned Ca^2+^ signals in WT from both ACC and BLA regions. **D.** Area under curve (AUC) of shock event-aligned Ca^2+^ signals in *Sh3^-/-^* from both ACC and BLA regions. **E-F**. Peak amplitude and frequency of spontaneous Ca^2+^ signals in WT observer ACC neurons and BLA neurons were compared. **G-H**. Peak amplitude and frequency of spontaneous Ca^2+^ signals in *Sh3^-/-^*observer ACC neurons and BLA neurons were compared. and *Sh3^-/-^* observer at HP and SP. Compared to HP, WT BLA spontaneous Ca^2+^ transients at SP showed increased peak amplitude but *Sh3^-/-^* showed no change. At baseline level, frequency was bigger in *Sh3^-/-^* BLA than WT. **I.** Schematic drawing showed inhibitory optogenetics manipulation on social fear transfer behavior in *Sh3^-/-^* mouse. eNpHR virus expression in ACC and BLA and fiber optic cannula implant in BLA were validated post-mortem. Scale bar is 500 μm. **K-N.** Compared to control group (*Sh3^-/-^*+YFP), inhibition on ACC to BLA projection (*Sh3^-/-^* +eNpHR) showed decreased freeze at SP and similar freeze at RP.

We first measured evoked Ca^2+^ spikes in neuron populations in the ACC and BLA of WT and *Sh3^-/-^* observers during the SP phase of social fear transfer. In ACC neurons, average area under curve (AUC) of shock-evoked Ca^2+^ spikes was highly correlated with average freeze duration in *Sh3^-/-^* and WT observers (**Fig. S4A-B**). In *Sh3^-/-^* observers, the AUC averaged across trials 1-12 was similar to WT however the AUC was significantly higher in *Sh3^-/-^*observers on average in trials 1-2 compared to WT (**Fig. S4C**). In contrast, in BLA neurons, average AUC of shock-evoked Ca^2+^ spikes in observers were highly correlated with average freeze duration in WT observers and not correlated in *Sh3^-/-^* observers (**Fig. S4D-E**). In *Sh3^-/-^* observers, the AUC averaged across trials 1-12 and the average of trials 1-2 was lower compared to WT observers (**Fig. S4F**).

To understand how ACC-BLA connectivity was involved in social fear transfer, we plotted shock-evoked Ca^2+^ spikes from 1^st^ to 12^th^ trial. In WT, analysis in ACC neurons of AUC of shock-evoked Ca^2+^ spikes showed invariant and non-adapting Ca^2+^ responses, while analysis in ACC neurons of AUC showed initial facilitation followed by a sustained plateau of Ca^2+^ responses (**Fig. 4C**). Strikingly, in *Sh3^-/-^*, analysis in both ACC and BLA neurons of AUC showed oscillatory non-monotonic Ca^2+^ responses (**Fig. 4D**). Different from WT in which Ca^2+^ responses in BLA neuron was significantly higher than the ones from ACC, *Sh3^-/-^* Ca^2+^ responses in BLA neuron was significantly lower than the ones from ACC (**Fig. 4C-D**). Taken together, the ACC-BLA circuit in *Sh3^-/-^* mice displayed curtailed and heterogeneous neuronal dynamics compared to the homeostatic synaptic transmission in WT mice. This suggests that ACC-BLA equilibrium maintenance responsible for appropriate social fear transfer response was disrupted in *Sh3^-/-^* mice.

Although we observed a general low correlation between average freeze duration and the amplitude or frequency of spontaneous Ca^2+^ spikes in both ACC and BLA neurons (**Fig. S4G-N**), there were detectable differences in Ca^2+^ activity changes from HP to SP between WT and *Sh3^-/-^*mice. In WT, BLA neurons presented higher average peak amplitude and lower frequency of Ca^2+^ spikes across both HP and SP phases compared to ACC neurons. However, in *Sh3^-/-^*, average peak amplitude of BLA neurons was similar to ACC neurons, which is different from WT. Moreover, comparing SP to HP phases, peak amplitude of BLA neurons increased in WT while it did not change in *Sh3^-/-^*. These differences of spontaneous Ca^2+^ spikes in BLA neurons suggest decreased activation of BLA neurons during social fear transfer in *Sh3^-/-^*.

To independently validate that ACC-BLA circuit dysfunction is causing increased social fear responses in *Sh3^-/-^*mice, we focally inhibited the ACC-BLA projection using an optogenetic approach. We injected virus carrying the inhibitory optogenetics opsin (eNpHR) or control opsin (YFP) into the ACC and implanted fiber optic cannula in the BLA in *Sh3^-/-^* observer mice which were then subjected to the 3-day social fear transfer paradigm (**Fig. 4I**). Optogenetic suppression of the ACC-BLA projection was achieved via yellow light stimulation (650nm wavelength) delivered to observers at the onset of SP. Optogenetic suppression of the ACC-BLA circuit decreased observational fear and did not alter fear retrieval compared to YFP controls (**Fig. 4K-N**). These results indicate that inhibition of the ACC projection to BLA curtailed hypersensitive fear transfer responses in *Sh3^-/-^*mice.

### *Sh3^-/-^* observer mouse displayed different social fear transfer-activated cFOS expression represented distinct neural network compared to WT mouse

To identify the neural network involved in heightened observational fear due to loss of *Shank3*, we measured cellular expression of cFOS, an immediate early gene that is rapidly and transiently expressed upon neural activity, combining iDisco whole brain clearing and cFOS immunohistochemistry. We measured cFOS expression across 341 brain regions of WT and *Sh3^-/-^* mice at 1.5 hours after completion of SP in the fear transfer test (transfer), and in control brains from WT and *Sh3^-/-^* mice which received no stimulation (homecage) (**Fig. 5A, Fig. S5A**). In brains from WT-transfer, cFOS was higher in the pallidum and thalamus, and lower in the primary somatosensory region, compared to brains from WT-homecage (**Fig. S5B**). It suggested sensory and motor control regions were associated with social fear transfer process. However, different social fear transfer brain network was revealed in brains of *Sh3^-/-^* mice. In brains from *Sh3^-/-^*-transfer, cFOS was increased in ACC and somatosensory region but decreased in thalamus (VENT), compared to brains from *Sh3^-/-^*-homecage (**Fig. S5C**). When comparing WT-transfer vs. *Sh3^-/-^*-transfer, increased cFOS was observed in ACC and somatosensory region but decreased cFOS was observed in pallidum. It suggested that ACC and pallidum are likely associated with exaggerated fear emotion response in *Sh3^-/-^* mice (**Fig. S5D**). Furthermore, cFOS was differentially expressed in striatum and thalamus regions in comparisons between WT-homecage and *Sh3^-/-^*-homecage (**Fig. S5E**). Taken together, these results identified substantially different anatomical networks between WT and *Sh3^-/-^* brains (**Fig. S5F**), including cortex, striatum, pallidum, thalamus and hippocampus regions, which may link the exaggerated fear emotion response to disrupted distinct brain circuit.

**Figure 5.**
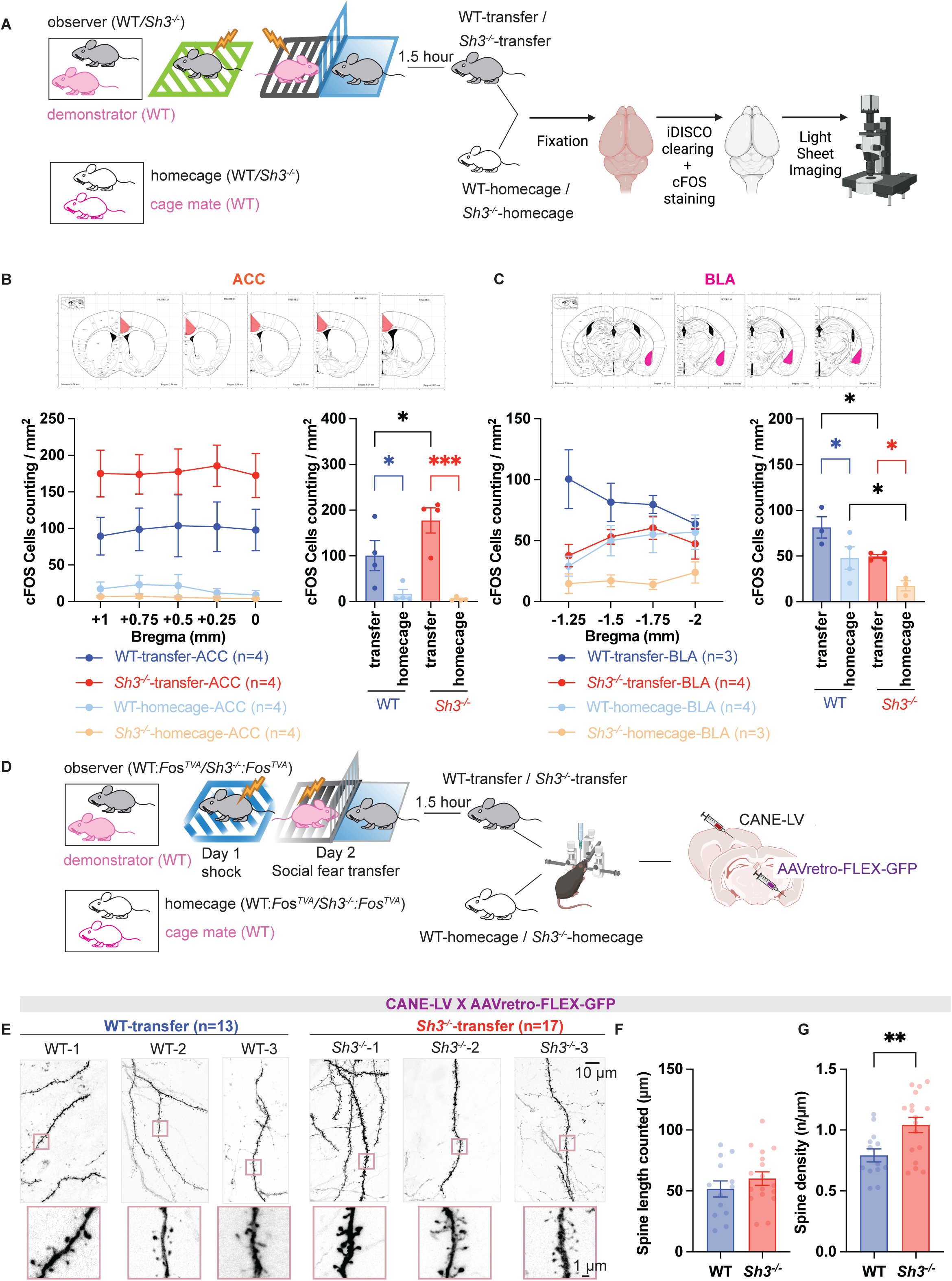
Sh3^-/-^ observer mouse heightened social fear transfer was encoded by hyperactive BLA-projecting ACC and hypoactive BLA ensembles. **A.** Schematic drawing showed that four groups of brains (WT-transfer, *Sh3^-/-^*-transfer, WT-homecage, *Sh3^-/-^*-homecage) were used for iDISCO brain clearing, cFOS immunofluorescence and light sheet microscopy imaging **B-C.** cFOS positive cells were counted from five different planes from ACC regions and four different planes from BLA regions. In ACC, both WT- and *Sh3^-/-^*-transfer groups showed significantly increased cFOS expression in response to social fear transfer than homecage groups. *Sh3^-/-^*-transfer increased cFOS expression than WT-transfer while *Sh3^-/-^*-homecage group showed similar cFOS expression as WT-homecage group (**B**). In BLA, both WT- and *Sh3^-/-^*-transfer groups showed significantly increased cFOS expression in response to social fear transfer than homecage groups. But *Sh3^-/-^* BLA showed lower cFOS expression than WT BLA at both transfer group comparison and homecage group comparison. **D.** Schematic drawing showed that BLA-projecting ACC neurons were labeled using CANE method. Injection of CANE-LV virus to ACC combining with injection of AAVretro-FLEX-GFP virus to BLA were performed in both WT and *Sh3^-/-^* mice, respectively. **E-F**. Example of GFP-expressed neuronal dendrites in ACC which were selective BLA-projecting ACC neurons and were strongly activated in transfer group. Spine density of GFP-labeled neurons was analyzed. Similar length of individual spine between WT and *Sh3^-/-^* was counted (**F**). BLA-projecting ACC neurons in *Sh3^-/-^* observer exhibited increased spine numbers than WT observer. (**G**)

To explore the contributions from ACC and BLA subregions, we measured cFOS expression at different Bregma locations. In the ACC region, at baseline, WT and *Sh3^-/-^* homecage control brains showed similar cFOS expression. Social fear transfer significantly increased cFOS expression in both WT and *Sh3^-/-^*brains compared to homecage controls. After social fear transfer, *Sh3^-/-^*brains showed increased cFOS expression in ACC regions from bregma +1mm to 0 mm compared to WT brains (**Fig. 5B**). In the BLA region, social fear transfer significantly increased cFOS expression in WT and *Sh3^-/-^* brains compared to homecage controls but cFOS activation was different between anterior and posterior BLA regions. *Sh3^-/-^*brains had lower cFOS expression than WT brains in BLA regions from bregma -1.25mm to -2 mm in homecage controls and after social fear transfer (**Fig. 5B**). These results indicate that hyperactive ACC and hypoactive BLA ensembles may underly heightened social fear transfer behavior in *Sh3^-/-^* mice.

To confirm that changes in circuitry of ACC projection to BLA contributed to aberrant social fear transfer behavior in *Sh3^-/-^* mice, we utilized captured activated neural ensembles (CANE) techniques to precisely target and mark transiently active BLA-projecting ACC neurons^66^ (**Fig. 5D, Fig. S6A**). First, we confirmed social fear transfer activated neuron ensembles capture in ACC and BLA by co-injecting EnvA^M21^ coated pseudotyped lentivirus carrying lenti-hSyn-Cre (CANE-LV) and AAV-FLEX-GFP virus in WT*:Fos^TVA^* and *Sh3^-/-^:Fos^TVA^*mice. Transfer groups showed abundant GFP labeled neurons in both ACC and BLA. In contrast, homecage group showed more GFP labeled neurons in BLA than ACC (**Fig. S6B**). Then we co-injected CANE-LV and AAV-FLEX-mCherry viruses in ACC and AAVretro-FLEX-GFP viruses in BLA to capture BLA-projecting ACC neuron ensembles. Transfer group captured significantly more GFP and mCherry labeled neurons than homcage group. GFP labeled neurons were overlapped with mCherry signals indicating successful capturing social fear transfer activated BLA-projecting ACC neurons (**Fig. S6C**). By analyzing high-resolution images of the dendrites of GFP-labeled neurons, we found that social fear transfer-captured BLA-projecting ACC neurons displayed higher spine density in *Sh3^-/-^*mice than WT mice (**Fig. 5E-G**), which might be associated with increased cFOS expression in ACC in *Sh3^-/-^*mice (**Fig. 5B**).

Overall, to understand how empathy changes in ASD, we evaluated the ASD mouse fear emotion responses to its social partner’s fear emotion using the social fear transfer paradigm and found that *Sh3^-/-^* mouse was hypersensitive to social fear transfer process than WT mouse. Combining molecular and circuitry manipulation tools, we demonstrated neural basis of social fear transfer processes in both WT and *Sh3^-/-^*mouse and established causal link between *Shank3* deficiency in ACC-BLA circuit and the abnormal empathic responses.

Significantly, our results for the first time contributed to understanding “double empathy” theory in ASD individuals mechanistically and established a framework for modeling empathy in ASD mouse models. Social fear transfer paradigm had distinct advantages in dissociating affective and cognitive components of empathy. *Sh3^-/-^* mice showed heightened affective responses but lacked cognitive component essential for retrieval of fear emotion. This conclusion extends recent finding showing *Sh3^-/-^*mice lack pain transfer responses^24^, which might be confounded by altered pain sensitivity in *Sh3^-/-^*mice and may reflect complexity of empathy representation in *Sh3^-/-^*mice.

Although we showed convincing results of heightened fear emotion in *Sh3^-/-^* mice, fear emotion and anxiety have some similarities in the accompanying physiological response and shared similar neural basis such as amygdala ^67,68^. Since anxiety is a common comorbidity of ASD and *Sh3^-/-^* mouse is known to exhibit elevated anxiety, endeavor to isolate anxiety from fear responses during social fear transfer was warranted in next. Particularly, we currently characterized weak responsive BLA neurons to social fear transfer in *Sh3^-/-^* mouse, but it would be interesting to know how BLA and other subregions of amygdala coordinates in mediating social fear transfer and whether such coordination is disrupted in *Sh3^-/-^*mouse. Understanding the neural circuits that mediate fear would also help to unravel the circuits of anxiety accompanied in ASD.

It has been reported that *Shank3* mutation mediated abnormal social behavior either through hypoactive ACC neurons with less spines ^56^, or hyperactive PFC neurons with more spines^57^. Our work not only added heightened affective empathy to the rich repository of social behavior abnormality of *Shank3* mutant mouse but also uncovered complicated dysfunctions of ACC regions. It suggested that characterizing different subtypes of ACC neuron in various social context would clarify how *Shank3* mutation impacts social behavior. ACC neuron axon efferent target to many brain regions and BLA is not the densest region. But ACC projection to BLA has been shown to play a critical role in integrating sensory and social stimuli, generating emotional presentation and executing decision-making in human and non-human animal brains ^69^. Disrupted equilibrium on ACC-BLA circuit in *Sh3^-/-^* mouse might not only mediate heightened social fear transfer. Assessing other social behavior domains using brain region specific manipulation tools would offer more insights on the function of ACC-BLA circuit.

Recent advancement on studying social behavior, emotion, and human genomics made it possible to model abnormal empathy in ASD mouse models. Utilizing cutting-edge techniques and interdisciplinary approaches to decompose empathy process in ASD, our work for the first time established a proof-of-concept to study interplay between gene and affective empathy in mouse, offered valuable insights on the social emotion process, and laid the groundwork for intervention for ASD and other neuropsychiatric disease.

However, there are a few limitations for this study: 1. Although we observed significant increased cFOS expression in ACC but increase of neural activity through *in vivo* Ca^2+^ recording was less impressive, it implicated that imbalance of inhibition and excitation might curtail the recorded neural activity in behavioral animals since our GCaMP was expressed in both inhibitory and excitatory neurons. Future study dissecting specific cell types is warranted. 2. BLA is a very complex region containing various subregions. Our cFOS analysis revealed differential results on anterior and posterior BLA indicating that BLA involvement on social fear transfer is subregion specific. Although our results implicated that neurons of ACC projection to BLA are hyperactive, we did not capture the increased activity for output neurons. Future study of examination of BLA subregions receiving ACC projection during social fear transfer will provide valuable mechanistic insights on how loss of *Shank3* leads to heightened affective empathy.

## Author contribution

Conceptualization: S-N.Q. and Y-H.J.

Methodology: S-N.Q. and Y-H.J.

Investigation: S-N.Q. and Y-H.J.

Visualization: S-N.Q.

Funding acquisition: S-N.Q. and Y-H.J.

Project administration: S-N.Q. and Y-H.J.

Supervision: S-N.Q. and Y-H.J.

Writing – original draft: S-N.Q.

Writing – review & editing: S-N.Q. and Y-H.J.

## Data and materials availability

All data are available in the main text or the supplementary materials. All materials used in the analysis are available to disseminate when requested.

---

**Fig. S1.**
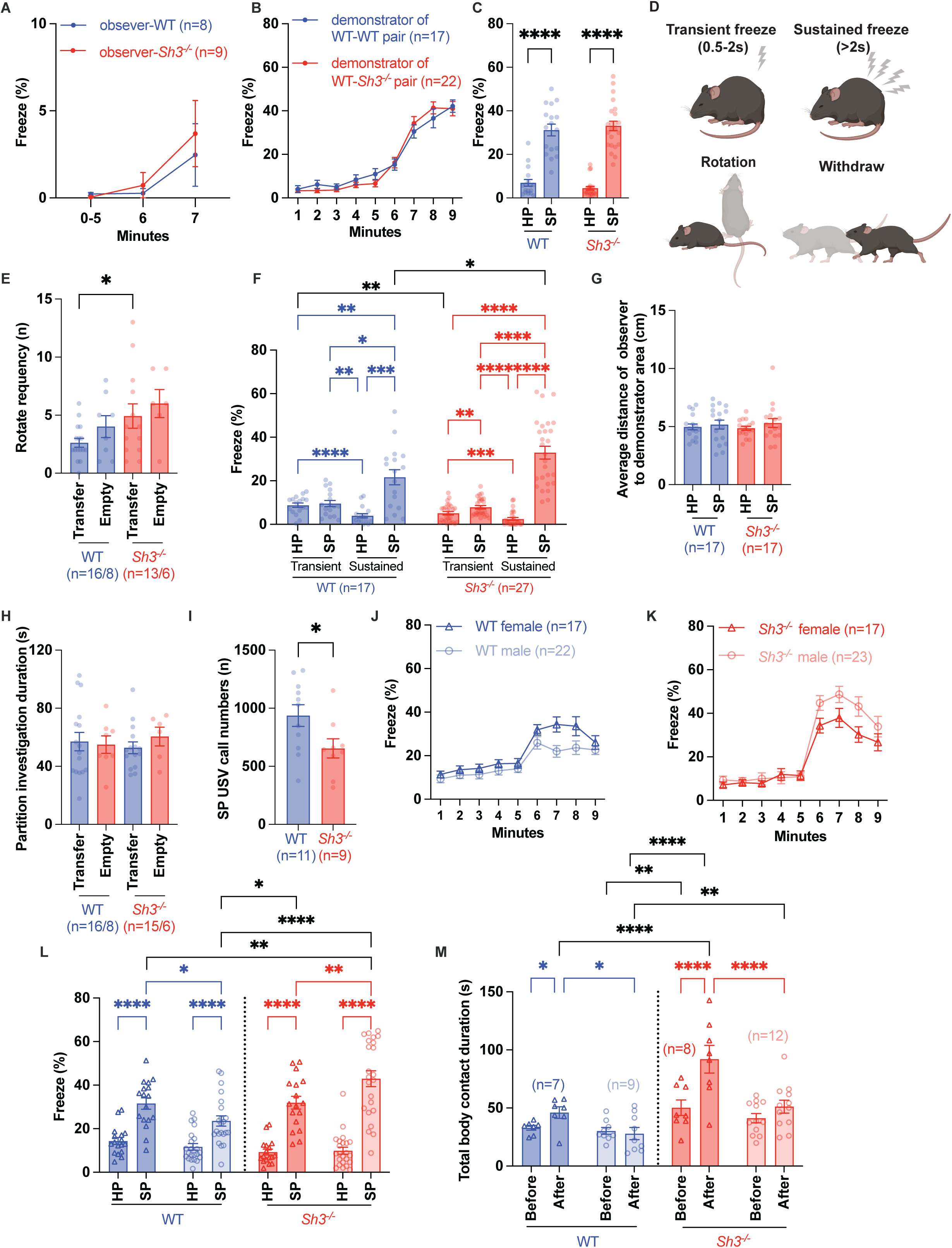
*Sh3^-/-^* observer mouse exhibited elevated fear emotional presentation. **A.** WT and *Sh3^-/-^* observer mouse freeze responses after shock were similar at day 1. After 5-minute habituation, two 2-second 0.3mA shocks were delivered at minute 5 and minute 6. No significant difference of freeze was found between WT and *Sh3^-/-^* observers. **B-C**. Freeze behavior of WT demonstrator mice from WT-WT and WT-*Sh3^-/-^* pairs at day 2 were similar. After 5-minute habituation, WT demonstrator significantly increased freeze when shocks began (**B**). On average, No significant difference of freeze was found between WT-WT pair and WT-*Sh3^-/-^* pair (**C**). **D-F.** Four types of fear behaviors, including transient freeze, sustained freeze, rotation and withdraw were compared (**D**). Although *Sh3^-/-^* observers exhibited higher rotating frequency than WT during transfer, no significant difference was found in both genotypes between transfer and empty condition (**E**). On average, both WT and *Sh3^-/-^* observers exhibited increased sustained SP freeze (sustained SP vs. transient SP, sustained SP vs. sustained HP, sustained SP vs. transient HP) and decreased sustained HP freeze (sustained HP vs. transient HP, sustained HP vs. transient SP). But compared to WT, *Sh3^-/-^* observer significantly increased sustained SP freeze (WT sustained SP vs. *Sh3^-/-^* sustained SP) and reduced transient HP freeze (WT transient HP vs. *Sh3^-/-^*transient HP) (**F**). **G-H**. WT and *Sh3^-/-^* observer mice showed similar exploratory behavior in the chamber. The distance between WT and *Sh3^-/-^* observers to the demonstrator area were similar at HP and SP, and WT and *Sh3^-/-^* observers spent similar time on actively investigating the partition either in presence of shocked demonstrator (transfer) or in absence of shocked demonstrator (empty). **I.** Less USV calls were accompanied with heightened social fear responses in *Sh3^-/-^* observers. During fear transfer, WT-WT pair made significant more USV calls than WT-*Sh3^-/-^* pair at SP. **J-L.** Average freeze responses of male (circle symbol) and female (triangle symbol) WT or *Sh3^+/-^* mice were compared at different time. **M.** Total body contact duration during free social interaction before and after social transfer process was compared between female and male pairs.

**Fig. S2.**
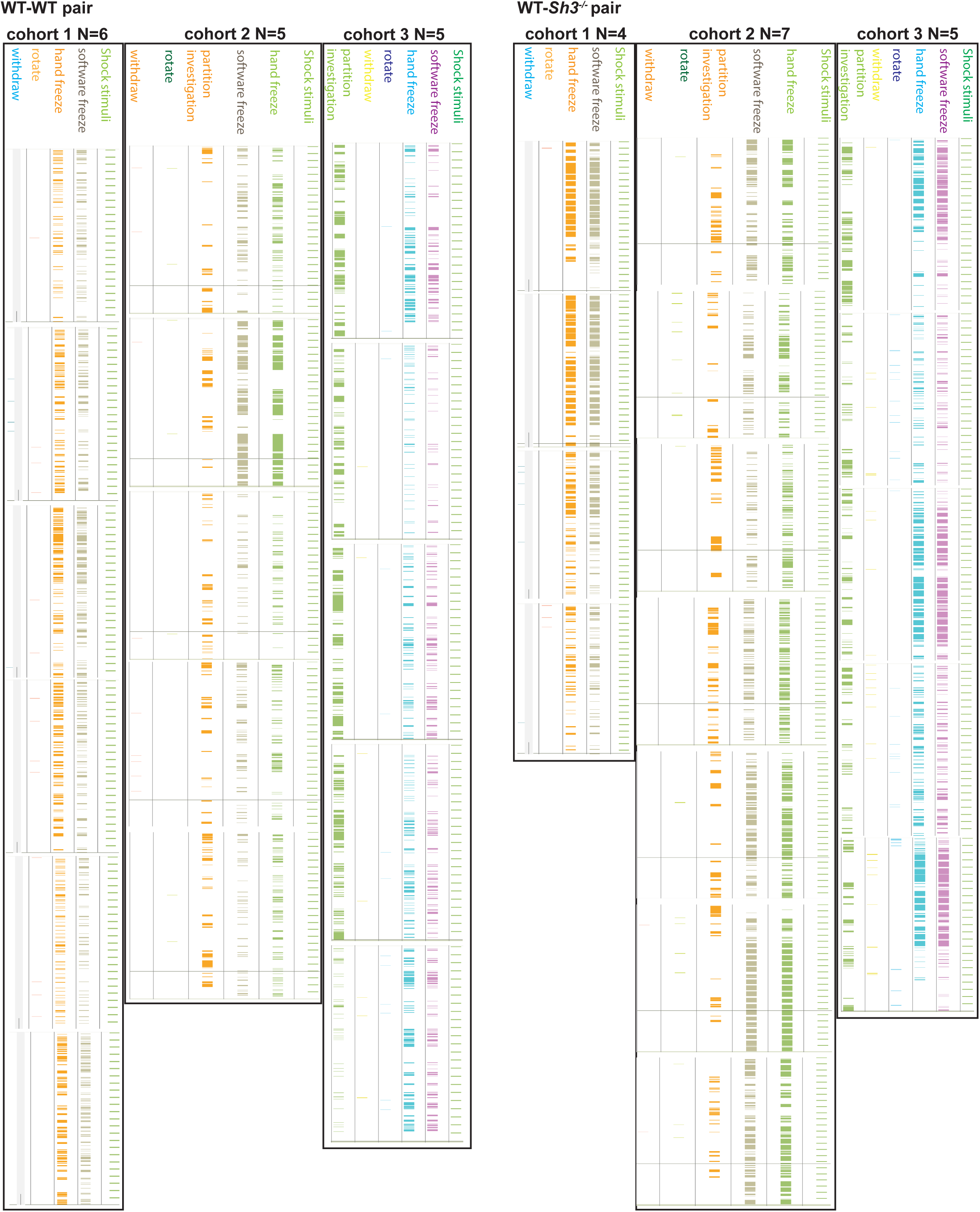
Colormap of individual WT and *Sh3^-/-^* observer mouse fear behaviors during fear transfer from. Detailed fear behaviors of observers from 16 WT-WT pairs and 16 WT*-Sh3^-/-^* pairs across three different cohorts were shown.

**Fig. S3.**
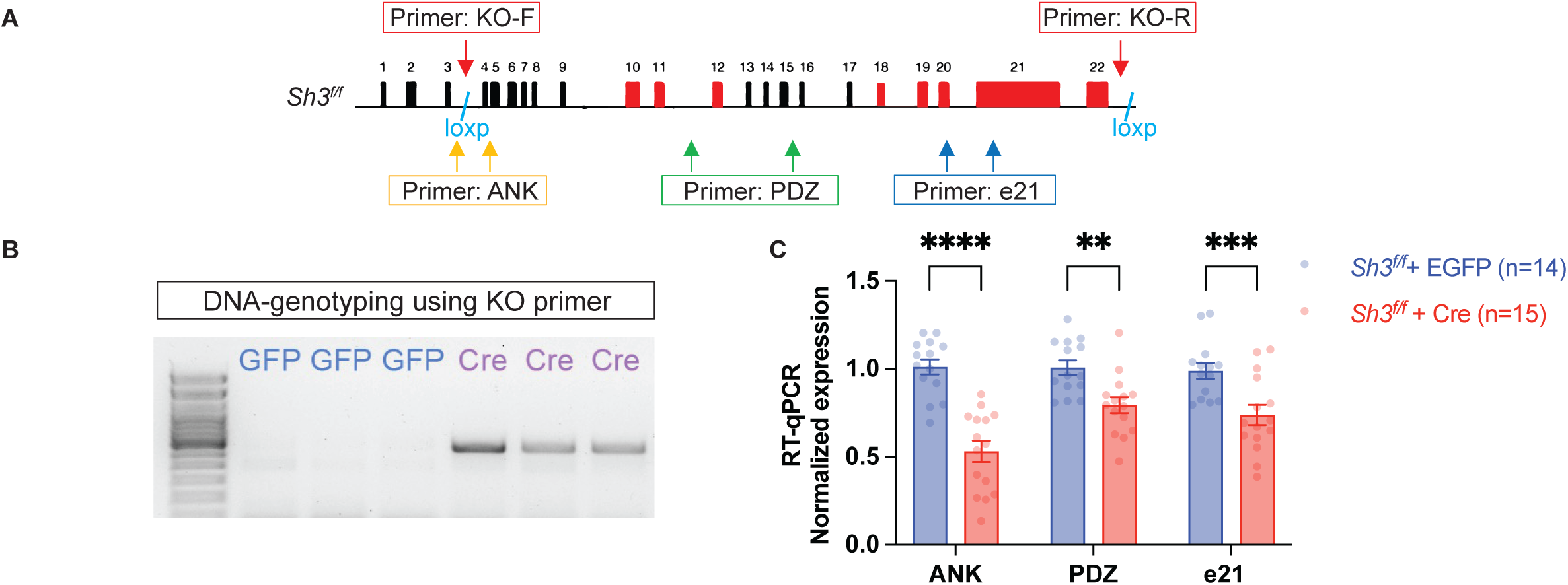
Molecular validation of *Shank3* knockdown in *Sh3^f/f^* mice. **A-C.** Primer-KO was designed for identifying 65kb *Shank3* DNA deletion between two loxp sites, which is located between exon 3 and 4, and after exon 22, respectively. Primer-ANK, Primer-PDZ and Primer-e21 were designed to evaluate *Shank3* RNA knockdown efficacy in different loci of *Shank3* gene (**A**). Genotyping of DNA from AAV-Cre-EGFP (Cre) or AAV-EGFP (GFP) viruses injected brain tissues showed that, Cre group presented amplification of a 500bp band but GFP group showed no PCR bands (**B**). RT-qPCR of RNA from virus injected brain tissues showed that Cre group showed reduction of *Shank3* RNA expression in various regions of *Shank3* gene(**C**).

**Fig. S4.**
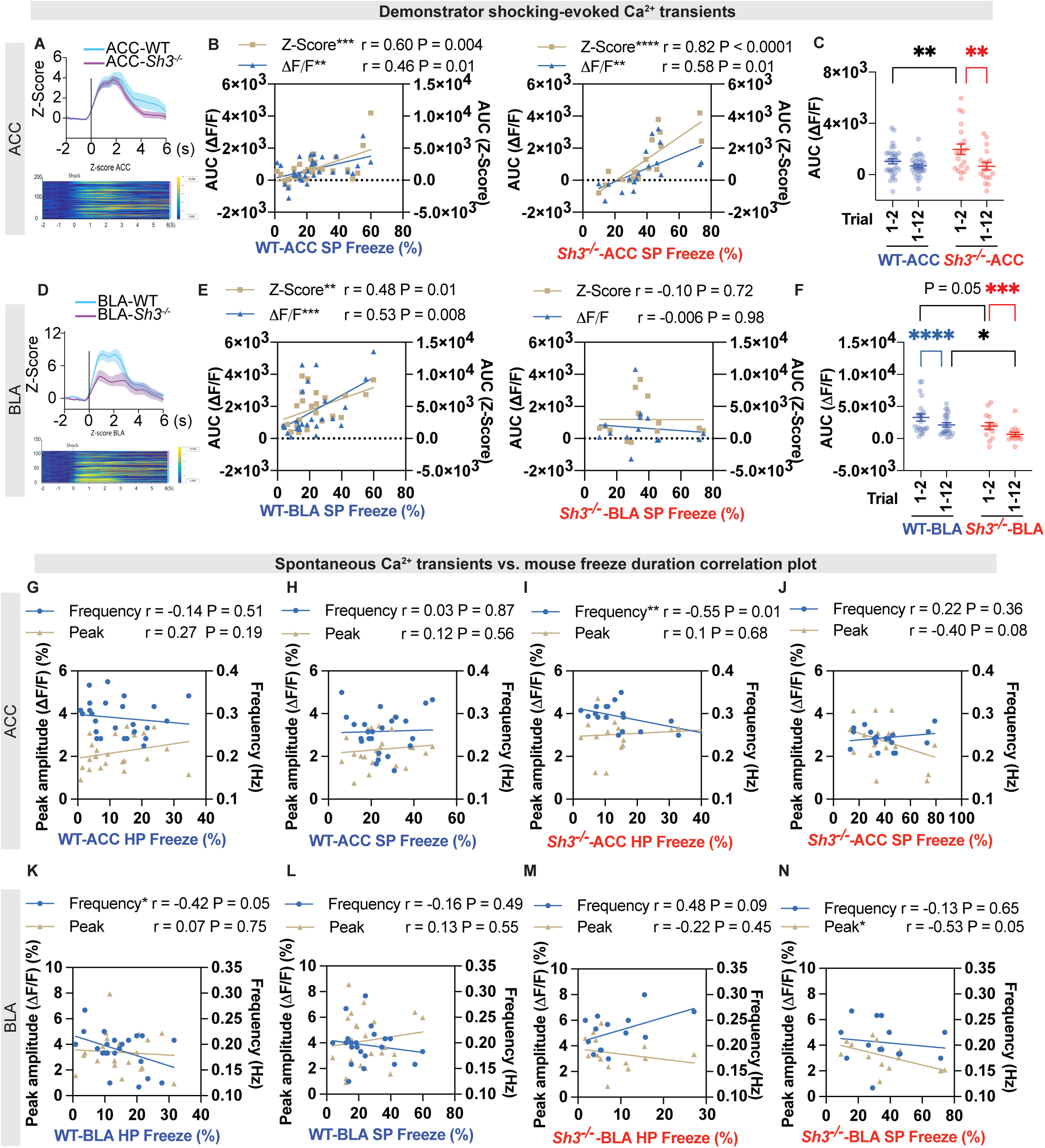
Comparison of spontaneous Ca^2+^ signals and demonstrator-shocking evoked Ca^2+^ signals from ACC and BLA between WT and *Sh3^-/-^* observer mouse. **A.** In ACC, Z-Score average of event-aligned Ca^2+^ signals from total 12 trials between WT and *Sh3^-/-^*observers was shown in line plot (top) and heatmap (bottom, each line represents one mouse). AUC of Ca^2+^ signals at 6-second post-stimuli was calculated with 2-second pre-stimuli as baseline. **B.** Significant positive correlation between ACC AUC (Z-Score or △F/F) and mouse SP freeze were shown in both WT and *Sh3^-/-^* observers. **C.** Average ACC AUC from first 2 trials (trial 1-2) and 12 trials (trial 1-12) were compared between WT and *Sh3^-/-^* observers. Although *Sh3^-/-^* observer exhibited increased AUC of trial 1-2 than WT, *Sh3^-/-^* observer showed adaptation of Ca^2+^ signals with decrease of AUC from trial 1-2 to trial 1-12. **D.** In BLA, Z-Score average of event-aligned Ca^2+^ signals from total 12 trials between WT and *Sh3^-/-^* observers was shown in line plot (top) and heatmap (bottom, each line represents one mouse). AUC of Ca^2+^ signals at 6-second post-stimuli was calculated with 2-second pre-stimuli as baseline. **E.** Significant positive correlation between BLA AUC (Z-Score or △F/F) and mouse SP freeze were shown in WT instead of *Sh3^-/-^* observer. **F.** Both WT and *Sh3^-/-^* observers showed adaptation of Ca^2+^ signals with decrease of BLA AUC from trial 1-2 to trial 1-12. *Sh3^-/-^* observer showed reduction of AUC than WT at both trial 1-2 and trial 1-12. **G-J.** Correlation plots of ACC spontaneous Ca^2+^ signals against observer mouse freeze were compared at both HP and SP stages between WT and *Sh3^-/-^* observers. No correlation between spontaneous Ca2+ signals and mouse freeze was found in WT mouse. In contrast, significant negative correlation between frequency and mouse HP freeze and strong negative correlation between peak amplitude and mouse SP freeze were found in *Sh3^-/-^* observers. **K-N.** Correlation plots of BLA spontaneous Ca^2+^ signals against observer mouse freeze were compared at both HP and SP stages between WT and *Sh3^-/-^* observers. At HP, the correlation between frequency and freeze was significantly negative in WT while it was positive in *Sh3^-/-^*mouse. At SP, no correlation between spontaneous Ca^2+^ signals and mouse freeze was found in WT mouse. In contrast, *Sh3^-/-^* mouse showed significant negative correlation between peak amplitude and mouse freeze.

**Fig. S5.**
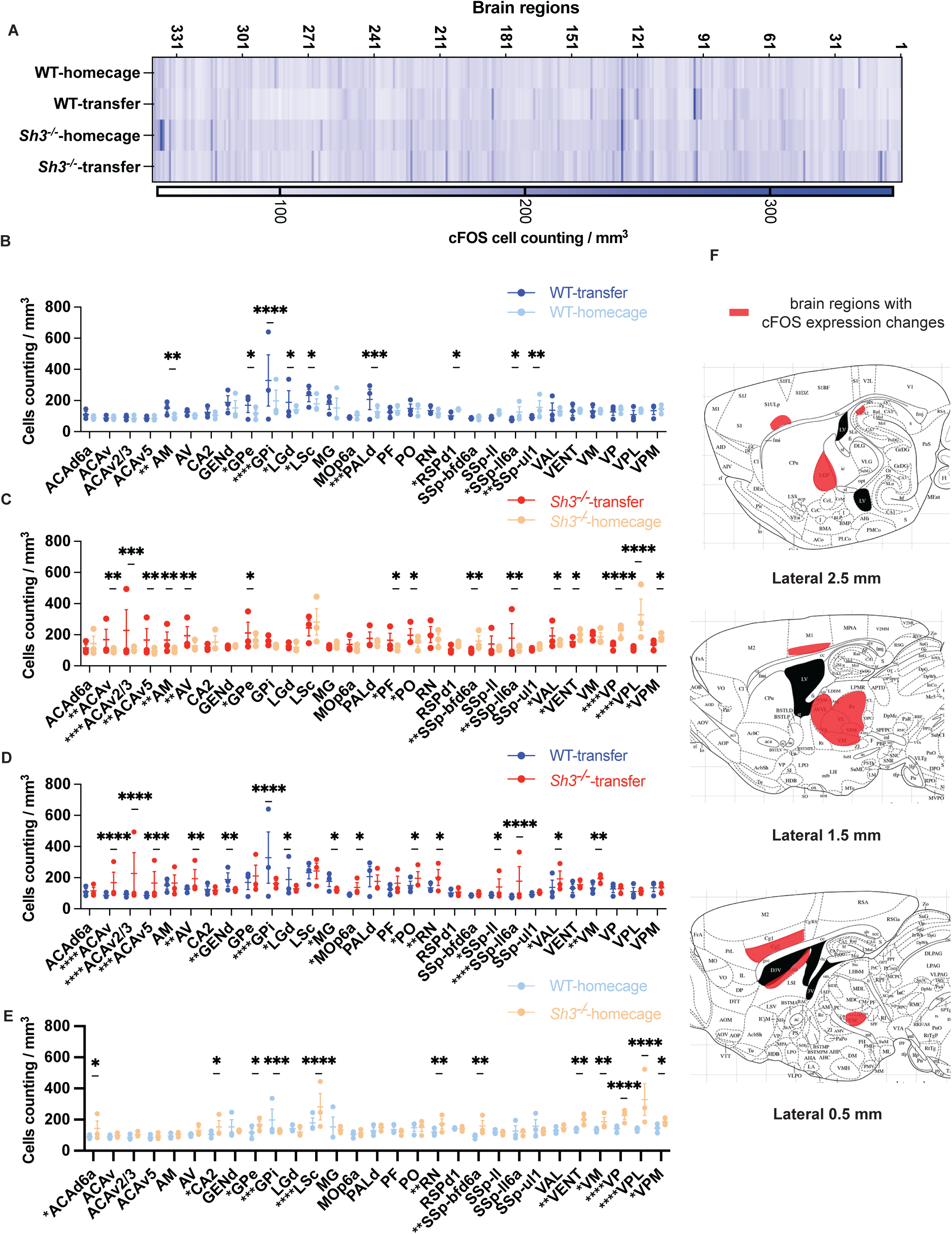
Whole brain cFOS quantification. **A.** Heatmap showed cFOS density from total of 341 brain regions across four groups of mice, WT-homecage, WT-transfer, *Sh3^-/-^*-homecage, and *Sh3^-/-^*-transfer. **B-F.** Four comparisons were analyzed using two-way ANOVA followed by post hoc multiple comparison: WT-transfer vs. WT-homcage (**B**), *Sh3^-/-^*-transfer vs. *Sh3^-/-^*-homecage (**C**), WT-transfer vs. *Sh3^-/-^*-transfer (**D**), *Sh3^-/^*-homecage vs. WT-homecage (**E**). Brain regions with significant change implicated in any of above comparisons were highlighted in the brain atlas as orange area (**F**). Olfactory bulb and cerebellum were cut off in atlas due to that those regions were damaged during imaging and results from those regions were not counted in the comparison.

**Fig. S6.**
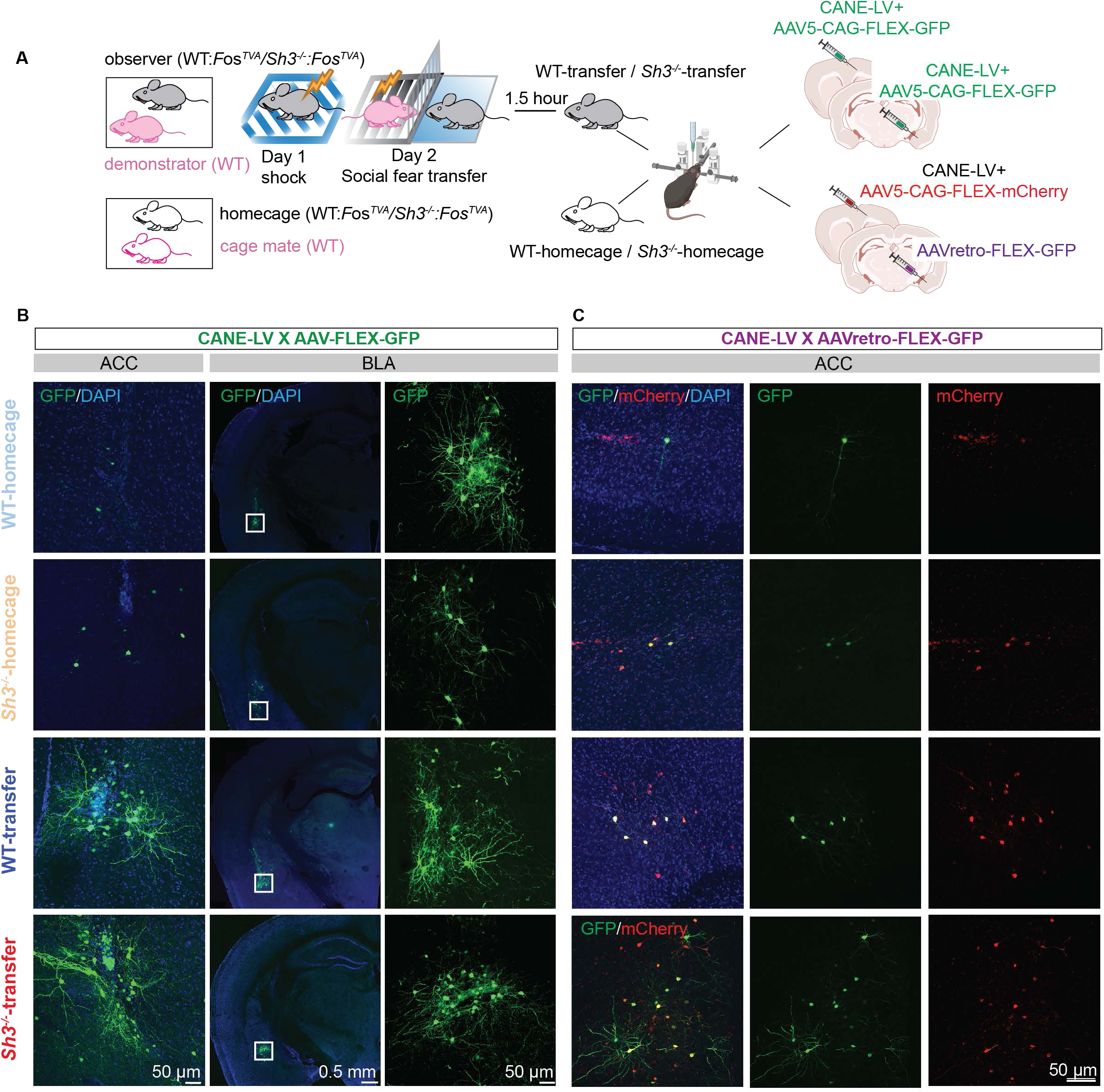
Captured social fear transfer activated ACC and BLA neurons in both WT and *Sh3^-/-^* mice using CANE method. **A.** Schematic drawing showed CANE experiments design and viruses used. **B.** Co-injection of CANE-LV and AAV-FLEX-GFP viruses were delivered to ACC or BLA in both WT and *Sh3^-/-^* mice, respectively. In ACC, transfer groups showed more GFP-expressed neurons while homecage group showed weak GFP expression in both WT and *Sh3^-/-^*. In contrast, in BLA, either transfer or homecage groups displayed abundant GFP-expressed cells in both WT and *Sh3^-/-^*BLA. **C.** Combining of CANE-LV and AAV-FLEX-mCherry viruses co-injection to ACC and AAVretro-FLEX-GFP injection to BLA was performed. GFP-signals were overlapped with mCherry labled cells, which were specifically BLA-projecting ACC neurons.

## Materials and Methods

### Animals

Animals were housed under standard 12 h/12 h light/dark cycles (7 AM-7 PM) with food, water ad libitum, controlled temperature and humidity in the Yale University animal facility. Experiments procedures conducted are following the Institutional Animal Care and Use Committee (IACUC) protocol at Yale University. *Shank3*^Δe4-22^ and *Shank3*^e4-22 flox/flox^ mouse lines were generated in-house and have been deposited to the Jackson Lab (#039524, #039525). *Fos^TVA^* mouse lines were obtained from Jackson Lab (#027831) and crossed with *Shank3*^Δe4-22^ mouse line for over 10 generations. Wild-type (WT), *Shank3*^Δe4-22^ homozygous (*Sh3^-/-^*) and *Shank3*^Δe4-22^ heterozygous mice (*Sh3^+/-^*) were used and obtained through *Sh3^+/-^* X *Sh3^+/-^* breeding. Wild-type *Fos^TVA^*homozygotes (WT*:Fos^TVA^*) and *Shank3*^Δe4-22^ homozygous *Fos^TVA^* homozygotes (*Sh3^-/-^:Fos^TVA^*) were used and obtained through *Shank3*^Δe4-22^ heterozygous *Fos^TVA^*homozygotes (*Sh3^+/-^:Fos^TVA^*) X *Sh3^+/-^:Fos^TVA^*breeding. *Shank3*^e4-22 flox/flox^ homozygous mice (*Sh3^f/f^*) were used and obtained through *Sh3^f/f^* X *Sh3^f/f^* breeding. At postnatal 21 days, all littermates of same cohort were weaned and genotyped then. Animals of the same sex were housed as two mice per cage, ie. demonstrator-observer dyads, until experiments day. Occasionally, after weaning, mice are housed as four mice per cage with two demonstrator mice and two observer mice to save cage space. Four-mice cage will be separated into 2 cages with demonstrator-observer dyad per cage at one week before experiments. Both male and female mice pairs were used. Genotype of demonstrator-observer dyad were as follows: WT-WT, WT-*Sh3^+/-^*, WT-*Sh3^-/-^*, WT-WT*:Fos^TVA^*, WT-*Sh3^-/-^:Fos^TVA^*, *Sh3^f/f^*-*Sh3^f/f^*. Two- to four-month-old mice were used. Mice with health problems such as runt, fight wounds, malocclusion, and eye problems, were excluded.

### Social fear transfer behavioral test

Mice were handled for 5 days before behavioral assays. Experiment procedures were performed within a time frame of 8 AM-6 PM. Behavioral tests were conducted once for each mouse. A soundproofed fear conditioning chamber (Med Associate) in Yale Rodent Behavior Analysis facility was used for day 1 shock experiment. A customized soundproofed fear transfer chamber (Ugo Basile) in a different dedicated procedure room was used for day 2 social fear transfer and day 3 social fear retrieval tests. Observer mouse and demonstrator mouse were littermate who were co-housed since weaning (over 30 days).

#### 3-day social fear transfer assay

At day 1, all cages of mice were transported from the housing room to the experiment room then mice were habituated in the room for 1 hour before tests. Only observer mouse from each cage was tested. After 5-minute habituation, two 2-second 0.3 mA electric shocks with interval of 1 minute were delivered to the observer. Experiment videos were recorded by a near infrared (NIR) camera and mouse immobility was scored by Med Associate freeze software. At day 2, individual mouse cage was transported from the housing room to procedure room one by one before each test. Observer mouse was put in the “observe section” and demonstrator mouse was put in the “shock section” of the fear transfer chamber. A customized transparent acrylic partitioner with 24 evenly spaced slits was put in the middle of the chamber enabling the sensory information exchange, including visual, auditory, olfactory, partial gustatory and tactile, between mouse pairs. At the “observe section” of the chamber, a transparent acrylic baseboard connecting with partition prevented observer mouse from getting shocked. At the “shock section” of the chamber, demonstrator mouse experienced electric shocks and a transparent acrylic lid connecting with partition prevented demonstrator mouse jumping into “observe section”. After 5-minute habituation, 24 2-second 0.7 mA electric shocks with interval of 10 seconds were delivered for 4 minutes. After then, the pair of mice were put back to the home cage. Then the cage was put back to the housing room and new cage of mouse pair was transported to the procedure room. Before new test, the chamber and the accessories were taken out and cleaned thoroughly. At day 3, the observer mouse was put back to the same chamber and 4-minute test was recorded. The behavioral video was recorded through a monochrome camera at the top of the fear transfer chamber. Mouse behavior metrics were analyzed using Ethovision XT15 software or manually scored. When analyzing mice fear behaviors, sustained freeze – immobility duration longer than two seconds, transient freeze – immobility duration between 0.5 to 2 seconds, rotation – rigid body rotation, and withdrawal – hind paw backward moving, were chosen according to previous studies^62,70^.

#### Stranger-pair social fear transfer assay

Two observer mice were co-housed and two demonstrator mice were co-housed in each cage. Observer mice and demonstrator mice came from different breeders and never housed in same cage. Similar to 3-day social fear transfer protocols, observer mouse experienced prior shocks at day 1, observed demonstrator mouse getting shocked at day 2 and put back to same chamber at day 3.

#### Naïve pair social fear transfer assay

Dyads of observer-demonstrator mice were co-housed since weaning. Observer mouse observed demonstrator mouse getting shocked at day 1 and put back to same chamber at day 2.

### Social dyad interaction behavioral test

Demonstrator-observer mouse pair social interaction is tested during 3-day social fear transfer assay. At day 2, right before fear transfer, the mouse pair was put in a Noldus Phenotyper chamber and freely interacted for 5 minutes. Then mouse pair experienced the 9-minute social fear transfer process. Right after that, the mouse pair was put back to same Noldus chamber and freely interacted for 5 minutes. The video was recorded by a camera at the top of the chamber. The body contact duration hand scored.

### Ultrasonic vocalization (USV) recording

USV recording protocol was as previously described^58^. In brief, an externally polarized condenser microphone with a frequency range of 10–200 kHz that was attached on top of “observer section” in the fear transfer chamber. The microphone was connected to the Avisoft-Ultrasound Gate recording software (Avisoft Bioacoustics). During the 9-minute social fear transfer process, the USV calls from the mouse pairs were recorded and save as WAV sound files using parameters optimized for mice. WAV files were converted to spectrograms and analyzed with automated whistle tracking parameters by the Avisoft SASLab Pro software. Because mouse pair rarely made calls during habituation period (first 5 minutes), only mouse calls during shocking period (5-9 minutes) were analyzed.

### Surgical procedure

After dyad mouse cage were set-up, Two- to three-month-old mice were used for surgery. Mice were anaesthetized by isoflurane vaporizer (RWD) and subcutaneous injected with analgesia buprenorphine (Ethiqa XR, 3.25 mg/kg), non-steroidal anti-inflammatory drug carprofen (5 mg/kg) and local anesthetic drug bupivicaine (5 mg/kg). Post-surgery care were done followed IACUC protocol.

### Optogenetics experiment

500nL AAV5-CaMKII-eNpHR-EYFP (UNC vector) or AAV5-CaMKII-EYFP (UNC vector) viruses were injected in ACC (AP +1mm, ML ±0.3mm, DV -0.45mm) bilaterally and two fiberoptic cannulas (Doric #MFC-200/250-0.66na-5.5mm) were implanted in BLA (AP -0.8mm, ML ±3.2mm, DV -4.95mm) bilaterally. The DV of cannula implant coordinate is 0.1 mm higher than that of virus injection. After recovering from surgery, observer mouse was single housed for 3 days to protect implant, during which period carprofen (5 mg/kg) were intraperitoneal injected daily for two days. Then observer mouse was co-housed with its original cage mate demonstrator mouse. If re-housed male mouse pairs caused fight and wounds, the mouse pairs were excluded. At 3-4 weeks after surgery, the mouse pairs went through the 3-day social fear transfer tests. Observer mice experienced two mild shocks at day 1. At day 2, a Doric optogenetics control unit was connected with cannula through a patch cord (Doric #SBP(2)_200/220/900-0.57na_2m_FCM-2xZF1.25(F)). Ethovision software controls the Doric optogenetics machine through TTL. Before test, observer mouse moved freely in the cage for 5 minutes for patch cord habituation. Then pairs of mice were put into the fear transfer chamber. After 5-minutes habituation in the chamber, continuous yellow light was delivered from optogenetics control unit to the mice during the 4-minute shock period. Then both mice were returned to home cage. At day 3, the observer mouse was put back to the same chamber for 4-minute recording. After tests, observer mice were perfused with 4% paraformaldehyde (PFA) and brains were collected and fixed with 4% PFA overnight. 100 µm thick brain slices were collected using vibratome (Leica VT200S) and imaged using confocal microscopy Zeiss LSM 900 to confirm that virus injection and cannula implant were correctly done. Mice with mistargeted virus injection or wrong cannula implant were excluded from data analysis.

### Capturing activated neuronal ensembles (CANE) experiment

For “transfer” group, pair of WT-WT*:Fos^TVA^*or WT-*Sh3^-/-^:Fos^TVA^* went through two mild shocks at day 1. For “homecage” group, pair of WT-WT*:Fos^TVA^* or WT-*Sh3^-/-^:Fos^TVA^* stayed in the cage. At day 2, “transfer” group observer mouse was put in stereotaxic surgery at 90 minutes after the fear transfer test, and “homecage” group observer mouse was put in stereotaxic surgery anytime at same day. Homemade CANE-LV viruses were generated as previously described^66,71^. In brief, HEK293T cells were transfected with the mixture construct of the pLenti-hSyn-Cre-WPRE (Addgene #86641), psPAX2 (Addgene #12260) and CANE-LV envelope plasmids (Addgene #86666). Virus media was extracted, concentrated and allocated as 5-10 µL vial stored in -80°C freezer. The titration of the CANE-LV viruses was determined as described^72^. Same titration of CANE-LV viruses (2.34 X 10^8^ TU/mL) was used for all experiments. To capture the social fear transfer-activated neurons, 500 nL mixture of 1:1 ratio of CANE-LV and AAV9-CAG-FLEX-EGFP viruses (Addgene #51502) was injected into right hemisphere ACC or right hemisphere BLA of observer mouse. To capture the BLA-projecting neurons in ACC, 500nL mixture of 1:1 ratio of CANE-LV and AAV5-EF1a-DIO-mCherry viruses (UNC vector) was injected into right hemisphere ACC combining with 500nL AAVretro-CAG-FLEX-EGFP virus (Addgene #51502) injected into right hemisphere BLA of observer mouse. After surgery, mouse was single housed for 3 weeks until when mouse brains were collected. After overnight 4% PFA fixation, 100 µm serial brain slices from Bregma AP +1.5mm to AP -2.5mm were collected using vibratome. All slices were examined using fluorescence microscopy and correct virus injected slices were imaged using Zeiss LSM900 confocal microscopy. For spine imaging, 63X oil objective len and 0.1 µm thickness was adopted for Z-stack imaging. At least 3 mice per genotype per group were used to confirm captured neurons in ACC and BLA and confirm the retrograde traced neurons from BLA to ACC.

### Fiber photometry experiment

Pairs of WT-WT and WT-*Sh3^-/-^*demonstrator-observer dyads were used for fiber photometry experiments. 500nL AAV5-hysn-GCaMP6s (UNC vector) virus was injected in one hemisphere ACC or BLA, respectively. After injection, the fiberoptic cannula (RWD #Black-200/250-0.37na-2mm) was implanted in ACC or the fiberoptic cannula (RWD #Black-200/250-0.37na-6mm) was implanted in BLA. The DV of cannula implant coordinate is 0.1 mm higher than that of virus injection. To protect the implant after surgery, the observer mouse was single housed for 3 days before co-housing with its original cage mate demonstrator mouse. In the case when male mice pairs fought and got wounds after co-housing, that male mice pairs were excluded from following experiments. At 3-4 weeks after surgery, the mouse pairs went through the 3-day social fear transfer tests. Observer mice experienced two mild shocks at day 1. At day 2, a patch cord (Doric #BBP(2)_200/220/900-0.37na_2m_FCM-2xZF1.25(F)_LAF) connected the cannula on mouse head with a fiber photometry system (RWD, R821 Tricolor Multichannel Fiber Photometry System). After putting on the patch cord, mouse moved freely in a clean cage for 5 minutes to adapt to the patch cord. Then both demonstrator and observer mouse were put to fear transfer chamber and *in vivo* Ca^2+^ signals were recorded simultaneously for 9-minute social fear transfer. After the behavior assays, observer mice were perfused with 4% PFA then brains were collected. 100 µm brain slices were collected and imaged to confirm the virus injection and cannula implant location were correct. Mistargeted virus injection or wrongly implanted cannula mouse data was excluded from data analysis.

#### Fiber photometry recording and analysis

A CMOS camera was used for synchronized acquisition at output of 470 nm and 410 nm LED. The 470 nm LED was used for GCaMP6s excitation and 410 nm LED was used as isosbestic control. LED power was ∼30 µW for 470 nm and ∼20 µW for 410 nm. Sampling frame rate is 30 fps. Start of recording and event of shock stimuli was synchronized and triggered by Ethovision software through TTL. RWD fiber photometry software was used for data acquisition and data analysis. Raw trace was preprocessed as follows: smoothing using moving average lowpass filter with coefficient W=10, baseline correction using iterative weighted partial least-squares (PLS) regression with coefficient β=5 and then motion correction using robust least-squares to fit 410 data to 470 to get motion-fitted410. Calculation formulas were as follows: △F/F = (F-F410)/F0, F = baseline corrected fluorescence data of 470 nm, F410 = baseline corrected fitted 410, F0 = median (baseline of raw data). Z-Score = (x-mean)/std, x = △F/F, mean = mean of △F/F, std = standard deviation of △F/F. Ca^2+^ peak statistics, including frequency and peak amplitude, obtained during habituation period (HP) data was from minute 2 to 4 of raw trace and during shocking period (SP) was from minute 5 to 7 of raw trace. Median absolute deviation (MAD) was first 0-5 second of the data with relatively stable baseline. The threshold of peak was calculated from the lowest point to highest point of the peak and was 2 to 3 folds of MAD. Minimum of time range of the peak was 1 second. To perform event analysis at SP, first 12 events with synchronized shock stimuli were chosen. Baseline of peri-event analysis was 2 second before event and area under curve (AUC) of Ca^2+^ transients during 6-second post time of event was analyzed. Total AUC of △F/F or Z-Score is the sum of area above and below reference of Y axis which is set at 0.

### Conditional knock-down *Shank3* experiment

Pairs of *Sh3^f/f^*-*Sh3^f/f^*were used. 500 nL AAV5-CMV-Cre-EGFP (Addgene #105545) or 500 nL AA5-CMV-EGFP (Addgene #105530) were injected into ACC or BLA bilaterally of 2-month-old *Sh3^f/f^* mice. At 3-4 weeks after injection, mice went through 3-day social fear transfer assays. After the behavior assays, observer mice brains were collected, and brain slices were cut and imaged to confirm the virus injection. Mouse with mistargeted virus injection was excluded from data analysis. Separate cohort of *Sh3^f/f^* mice were used for examining viral-induced *Shank3* knock-down efficiency. 500 nL AAV5-CMV-Cre-EGFP was injected into ACC or BLA in one hemisphere, then 500 nL AAV5-CMV-EGFP was injected into ACC or BLA in another hemisphere. At 3-4 weeks after injection, ACC and BLA brain regions in both hemispheres were collected for DNA and RNA extraction and RT-qPCR. DNA and RNA was isolated from the forebrain hemisphere using Trizol (Invitrogen, 15596018) followed manufacturer protocol. DNA was used for PCR to confirm the genotype of the brain tissue. Primer used for DNA genotyping is: *Shank3* KO forward, TTGCATCTGGGACCTACTCC, reverse, AAAGCACTGACTCCTCTCTTGG. Toal RNA treated with DNase I (Invitrogen, AM1907) to remove contaminating DNA was transcribed into cDNA and quantitative PCR was performed. Gene expressions were normalized to the expression of GAPDH. The primers used to amplify cDNAs are: *Shank3* ANK, forward, CAAGCGGAGAGTTTATGCCCAG, reverse, CCACCTTATCTGTGCTGTGTAGC; *Shank3* PDZ, forward, GTGGAAGAAGTGCAGATGCG, reverse, GCTGTCATAGGAA CCCACAG; *Shank3* e21, forward, TCATTGTTTGAGCGCCAGG, reverse, AGTAGGGAT GCCAGCTTCTC; *Gapdh* forward, CAAAATGGTGAAGGTCGGTG, reverse, AATG AAGGGGTCGTTGATGG.

### iDisco-cleared brain cFOS staining experiment

3-day social fear transfer behavior paradigm was used for iDisco brain clearing and cFOS staining experiments. For “transfer” group, pair of WT-WT or WT-*Sh3^-/-^* went through two mild shocks at day 1. For “homecage” group, pair of WT-WT or WT-*Sh3^-/-^*stayed in the cage. At day 2, “transfer” group observer mouse brain was collected via transcardiac 4% PFA perfusion at 90 minutes after the fear transfer test, and “homecage” group observer mouse brain was collected anytime at same day. After overnight 4% PFA fixation, brains were put in 30% sucrose for 2 weeks until iDisco clearing experiments. Half brains went through the iDisco brain clearing protocol as previously described^73^. Samples went through delipidation through being washed with B1n buffer (0.2 M Glycine, 0.1% (v/v) Triton X-100, 1 mM NaOH, 0.02% NaN_3_), dehydrated with series of methanol (20%, 40%, 60%, 80% methanol in B1n, 33% methanol in dichloromethane (DCM) (Sigma #270997) and 100% methanol), overnight bleach in 5% (v/v) H_2_O_2_ in methanol, rehydrated with series of methanol (80%, 60%, 40% and 20% methanol in B1n), and finally washed in PBS/0.2% (v/v) Triton X-100 for 1 hour twice, before further cFOS staining procedures. Staining was done in a 37 °C hybridization oven with a rotator. Samples were permeabilized in buffer containing 0.2% Triton X-100 /0.3 M glycine/ 20% DMSO/ 0.01% NaN_3_ overnight, blocked in buffer containing 0.2% Triton X-100 /6% donkey Serum/10% DMSO/0.01% NaN_3_ overnight, incubated in primary antibody anti-Rabbit cFOS (1:1000, CST#2250S) for 5 days, washed in PBS TwH buffer (0.2% Tween20/10 μg/ml heparin/0.01% NaN_3_) overnight, incubated in secondary antibody donkey-anti-rabbit Alexa Fluor 647 (1:2000, Invitrogen#A-31573) for 4 days, and washed in PBS overnight. Labeled brain samples were cleared in series of methanol (20%, 40%, 60%, 80% methanol in B1n, 33% methanol in DCM), 100% DCM, then sunk in dibenzyl ether (DBE) (Sigma #108014) solution and were ready for imaging.

First, samples were positioned in the coronal orientation and imaged using a 1.3x/0.08/WD6 LVMI-Fluor objective len in an Ultramicroscope light sheet microcopy (LaVision Biotec) equipped with a sCMOS camera (Andor Neo). 488-nm laser (emission filter 525/50nm) and 638-nm laser (emission filter 670/50nm) were used for dual light sheets imaging (sheet NA 0.147, sheet width 40%). Each channel was imaged independently under identical laser power and exposure settings across all samples. The voxel size of images was 5 × 2.5 × 2.5 μm³ (zyx). Imaris (Oxford Instruments) and Neuroinfo (MBF Bioscience) softwares were used for 3D images manipulation and alignment with Allen Reference Brain. Then ROIs of ACC and BLA regions were drawn and cFOS positive cells were counted using “Analyze particle” function with same threshold across all samples in the ImageJ software. ACC ROI containing all Anterior Cingulate Area (ACA) regions, and BLA ROI containing Lateral Amygdalar Nucleus (LA), Basolateral Amygdalar Nucleus anterior part (BLAa) and posterior part (BLAp) from Allen Reference Brain Atlas. 5 planes of images corresponding to Bregma +1 mm, +0.75 mm, +0.5 mm, +0.25 mm, and 0 mm were chosen for ACC quantification. 4 planes of images corresponding to Bregma -1.25 mm, -1.5 mm, -1.75 mm and -2 mm were chosen for BLA quantification.

To obtain high-resolution images for whole brain cFOS quantification, the same brain samples were sent to *Translucence Biosystems* for imaging and analysis. A ZEISS Lightsheet Z.1 microscope integrated with Translucence’s Mesoscale Imaging System was used for imaging. Samples were positioned in the sagittal orientation using custom sample holders and imaged with a 2.5x/0.12 NA detection optic at 1.0x optical zoom and 5x/0.1 NA illumination optics. Illumination was achieved with 488-nm and 638-nm lasers, paired with BP 505-545 and LP 660 emission filters, respectively. Each channel was imaged independently under identical laser power and exposure settings across samples. The voxel size of images was 5.84 × 1.82 × 1.82 μm³ (zyx). The czi output images were tiled and stitched using customized Stitchy software to generate tiff stacks of brain volumes. The tiff stacks were processed using customized Voxels software, which employs AI-enabled segmentation for tissue volumes. 3D probability maps and deterministic filters were used to segment individual cFOS positive neurons across the brain. Each brain was warped onto the Allen Reference Brain, and the warping parameters were applied to the coordinates of all identified neurons. This allowed for regional analysis of cFOS counts and signal intensities, based on the Allen Reference Atlas. To localize segmented neurons to brain regions, we transformed each brain into a common coordinate framework using the Allen Reference Brain and its anatomical atlas. Due to variations in anatomy (e.g., dissection, fixation, or clearing artifacts), the brains were warped to the reference atlas using the green autofluorescence channel from the light sheet images. Autofluorescence and signal channels were down sampled. Each sample was then warped to the atlas, and the signal channel was integrated with chromatic aberration correction. To ensure accurate registration, we performed rigorous quality control (QC) by comparing the x, y, and z coordinates of key anatomical landmarks in the warped images to the reference brain. Because precision is limited for very small regions and can be affected by damaged brain areas (e.g., cerebellum, olfactory bulbs), cerebellum and olfactory bulb regions, and regions’ volume smaller than 1000 μm³ were excluded from statistics comparison.

### Statistical analysis

Graphpad Prism 10.0 (GraphPad Software) was used for the statistical analysis and data plot. Differences were analyzed by student’s *t*-test if comparing two variables or using ANOVA if comparing more than two variables followed by multiple comparisons either using statistical hypothesis testing with correction or planned comparisons without correction. Significance was determined by two-tailed *t*-test *p* value or multiplicity adjusted *p* value. n represents the total number of data points per group, usually meaning the mice number with exception that n represents number of dendrites in spine density comparison. Both sexes of mice were used and balanced across genotype and treatment. Behavioral difference between male and female mice were explored. Individual data points were plotted and presented as average ± SEM. *p < 0.05* was considered statistically significant. Adobe Illustrator was used to create schematic diagram.

## Reference

1 Puścian, A. et al. Ability to share emotions of others as a foundation of social learning. Neuroscience & Biobehavioral Reviews 132, 23–36 (2022). 10.1016/j.neubiorev.2021.11.022

2 de Waal, F. B. The antiquity of empathy. Science 336, 874–876 (2012). 10.1126/science.1220999

3 de Waal, F. B. M. Empathy, the umbrella term. Neurosci Biobehav Rev 129, 180–181 (2021). 10.1016/j.neubiorev.2021.07.034

4 Preston, S. D. & de Waal, F. B. Empathy: Its ultimate and proximate bases. Behav Brain Sci 25, 1–20; discussion 20-71 (2002). 10.1017/s0140525x02000018

5 Preston, S. D. & Hofelich, A. J. The Many Faces of Empathy: Parsing Empathic Phenomena through a Proximate, Dynamic-Systems View of Representing the Other in the Self. Emotion Review 4, 24–33 (2012). 10.1177/1754073911421378

6 de Waal, F. B. M. & Preston, S. D. Mammalian empathy: behavioural manifestations and neural basis. Nat Rev Neurosci 18, 498–509 (2017). 10.1038/nrn.2017.72

7 Kanner, L. Autistic disturbances of affective contact. Nervous child 2, 217–250 (1943).

8 Fletcher-Watson, S. & Bird, G. Autism and empathy: What are the real links? Autism 24, 3–6 (2020). 10.1177/1362361319883506

9 Shah, P., Livingston, L. A., Callan, M. J. & Player, L. Trait Autism is a Better Predictor of Empathy than Alexithymia. J Autism Dev Disord 49, 3956–3964 (2019). 10.1007/s10803-019-04080-3

10 Stroth, S. et al. Empathy in Females With Autism Spectrum Disorder. Front Psychiatry 10, 428 (2019). 10.3389/fpsyt.2019.00428

11 Shi, L. J. et al. Altered empathy-related resting-state functional connectivity in adolescents with early-onset schizophrenia and autism spectrum disorders. Asian J Psychiatr 53, 102167 (2020). 10.1016/j.ajp.2020.102167

12 Shalev, I. et al. Reexamining empathy in autism: Empathic disequilibrium as a novel predictor of autism diagnosis and autistic traits. Autism Res 15, 1917–1928 (2022). 10.1002/aur.2794

13 Sucksmith, E., Allison, C., Baron-Cohen, S., Chakrabarti, B. & Hoekstra, R. A. Empathy and emotion recognition in people with autism, first-degree relatives, and controls. Neuropsychologia 51, 98–105 (2013). 10.1016/j.neuropsychologia.2012.11.013

14 Lord, C. et al. The Lancet Commission on the future of care and clinical research in autism. The Lancet 399, 271–334 (2022). 10.1016/S0140-6736(21)01541-5

15 Sun, W. et al. Reviving-like prosocial behavior in response to unconscious or dead conspecifics in rodents. Science 387, eadq2677 (2025). doi:10.1126/science.adq2677

16 Sheeran, W. M. & Donaldson, Z. R. An innate drive to save a life. Science 387, 827–828 (2025). doi:10.1126/science.adv3731

17 Choi, J. et al. Cortical representations of affective pain shape empathic fear in male mice. Nature Communications 16, 1937 (2025). 10.1038/s41467-025-57230-w

18 Zhang, M., Wu, Y. E., Jiang, M. & Hong, W. Cortical regulation of helping behaviour towards others in pain. Nature 626, 136–144 (2024). 10.1038/s41586-023-06973-x

19 Kim, S. W. et al. Hemispherically lateralized rhythmic oscillations in the cingulate-amygdala circuit drive affective empathy in mice. Neuron 111, 418–429 e414 (2023). 10.1016/j.neuron.2022.11.001

20 Zhang, M.-M. et al. Glutamatergic synapses from the insular cortex to the basolateral amygdala encode observational pain. Neuron 110, 1993–2008.e1996 (2022). 10.1016/j.neuron.2022.03.030

21 Terranova, J. I. et al. Hippocampal-amygdala memory circuits govern experience-dependent observational fear. Neuron 110, 1416–1431 e1413 (2022). 10.1016/j.neuron.2022.01.019

22 Takayama, K. et al. Autism Spectrum Disorder Model Mice Induced by Prenatal Exposure to Valproic Acid Exhibit Enhanced Empathy-Like Behavior via Oxytocinergic Signaling. Biol Pharm Bull 45, 1124–1132 (2022). 10.1248/bpb.b22-00200

23 Gonzales-Rojas, R. et al. The mouse model of fragile X syndrome exhibits deficits in contagious itch behavior. Sci Rep 10, 17679 (2020). 10.1038/s41598-020-72891-x

24 Rein, B. et al. MDMA enhances empathy-like behaviors in mice via 5-HT release in the nucleus accumbens. Science Advances 10, eadl6554 (2024). doi:10.1126/sciadv.adl6554

25 Allsop, S. A. et al. Corticoamygdala Transfer of Socially Derived Information Gates Observational Learning. Cell 173, 1329–1342 e1318 (2018). 10.1016/j.cell.2018.04.004

26 Burkett, J. P. et al. Oxytocin-dependent consolation behavior in rodents. Science 351, 375–378 (2016). 10.1126/science.aac4785

27 Smith, M. L., Asada, N. & Malenka, R. C. Anterior cingulate inputs to nucleus accumbens control the social transfer of pain and analgesia. Science 371, 153–159 (2021). 10.1126/science.abe3040

28 Jeon, D. et al. Observational fear learning involves affective pain system and Cav1.2 Ca2+ channels in ACC. Nat Neurosci 13, 482–488 (2010). 10.1038/nn.2504

29 Gu, X. et al. Anterior insular cortex is necessary for empathetic pain perception. Brain 135, 2726–2735 (2012). 10.1093/brain/aws199

30 Singer, T. et al. Empathy for pain involves the affective but not sensory components of pain. Science 303, 1157–1162 (2004). 10.1126/science.1093535

31 Huang, Z. et al. Ventromedial prefrontal neurons represent self-states shaped by vicarious fear in male mice. Nat Commun 14, 3458 (2023). 10.1038/s41467-023-39081-5

32 Ito, W. & Morozov, A. Prefrontal-amygdala plasticity enabled by observational fear. Neuropsychopharmacology 44, 1778–1787 (2019). 10.1038/s41386-019-0342-7

33 Mindaye, S. A., et al. Separate anterior paraventricular thalamus projections differentially regulate sensory and affective aspects of pain. Cell Reports 43 (2024). 10.1016/j.celrep.2024.114946

34 Bernhardt, B. C. & Singer, T. The neural basis of empathy. Annu Rev Neurosci 35, 1–23 (2012). 10.1146/annurev-neuro-062111-150536

35 Engen, H. G. & Singer, T. Empathy circuits. Curr Opin Neurobiol 23, 275–282 (2013). 10.1016/j.conb.2012.11.003

36 Kolevzon, A. et al. Phelan-McDermid syndrome: a review of the literature and practice parameters for medical assessment and monitoring. J Neurodev Disord 6, 39 (2014). 10.1186/1866-1955-6-39

37 Tabet, A.-C. et al. A framework to identify contributing genes in patients with Phelan-McDermid syndrome. npj Genomic Medicine 2, 32 (2017). 10.1038/s41525-017-0035-2

38 Monteiro, P. & Feng, G. SHANK proteins: roles at the synapse and in autism spectrum disorder. Nat Rev Neurosci 18, 147–157 (2017). 10.1038/nrn.2016.183

39 Li, J. et al. Spatiotemporal profile of postsynaptic interactomes integrates components of complex brain disorders. Nature Neuroscience 20, 1150–1161 (2017). 10.1038/nn.4594

40 Jia, B. et al. Shank3 oligomerization governs material properties of the postsynaptic density condensate and synaptic plasticity. Cell (2025). 10.1016/j.cell.2025.07.047

41 Bozdagi, O. et al. Haploinsufficiency of the autism-associated Shank3 gene leads to deficits in synaptic function, social interaction, and social communication. Molecular Autism 1, 15 (2010). 10.1186/2040-2392-1-15

42 Wang, X. et al. Synaptic dysfunction and abnormal behaviors in mice lacking major isoforms of Shank3. Hum Mol Genet 20, 3093–3108 (2011). 10.1093/hmg/ddr212

43 Peça, J. et al. Shank3 mutant mice display autistic-like behaviours and striatal dysfunction. Nature 472, 437–442 (2011). 10.1038/nature09965

44 Schmeisser, M. J. et al. Autistic-like behaviours and hyperactivity in mice lacking ProSAP1/Shank2. Nature 486, 256–260 (2012). 10.1038/nature11015

45 Speed, H. E. et al. Autism-Associated Insertion Mutation (InsG) of Shank3 Exon 21 Causes Impaired Synaptic Transmission and Behavioral Deficits. The Journal of Neuroscience 35, 9648–9665 (2015). 10.1523/jneurosci.3125-14.2015

46 Jaramillo, T. C. et al. Altered Striatal Synaptic Function and Abnormal Behaviour in Shank3 Exon4-9 Deletion Mouse Model of Autism. Autism Res 9, 350–375 (2016). 10.1002/aur.1529

47 Mei, Y. et al. Adult restoration of Shank3 expression rescues selective autistic-like phenotypes. Nature 530, 481–484 (2016). 10.1038/nature16971

48 Zhou, Y. et al. Mice with Shank3 Mutations Associated with ASD and Schizophrenia Display Both Shared and Distinct Defects. Neuron 89, 147–162 (2016). 10.1016/j.neuron.2015.11.023

49 Jaramillo, T. C. et al. Novel Shank3 mutant exhibits behaviors with face validity for autism and altered striatal and hippocampal function. Autism Research 10, 42–65 (2017). 10.1002/aur.1664

50 Harony-Nicolas, H. et al. Oxytocin improves behavioral and electrophysiological deficits in a novel Shank3-deficient rat. eLife 6, e18904 (2017). 10.7554/eLife.18904

51 Yoo, T. et al. GABA Neuronal Deletion of Shank3 Exons 14–16 in Mice Suppresses Striatal Excitatory Synaptic Input and Induces Social and Locomotor Abnormalities. Frontiers in Cellular Neuroscience 12 (2018). 10.3389/fncel.2018.00341

52 Yoo, Y.-E. et al. Shank3 Mice Carrying the Human Q321R Mutation Display Enhanced Self-Grooming, Abnormal Electroencephalogram Patterns, and Suppressed Neuronal Excitability and Seizure Susceptibility. Frontiers in Molecular Neuroscience 12 (2019). 10.3389/fnmol.2019.00155

53 Wang, L. et al. An autism-linked missense mutation in SHANK3 reveals the modularity of Shank3 function. Molecular Psychiatry 25, 2534–2555 (2020). 10.1038/s41380-018-0324-x

54 Chung, M. et al. Conditional knockout of Shank3 in the ventral CA1 by quantitative in vivo genome-editing impairs social memory in mice. Nat Commun 15, 4531 (2024). 10.1038/s41467-024-48430-x

55 Lee, D. K. et al. Reduced sociability and social agency encoding in adult Shank3-mutant mice are restored through gene re-expression in real time. Nat Neurosci 24, 1243–1255 (2021). 10.1038/s41593-021-00888-4

56 Guo, B. et al. Anterior cingulate cortex dysfunction underlies social deficits in Shank3 mutant mice. Nature Neuroscience 22, 1223–1234 (2019). 10.1038/s41593-019-0445-9

57 Kim, S. et al. Neural circuit pathology driven by Shank3 mutation disrupts social behaviors. Cell Rep 39, 110906 (2022). 10.1016/j.celrep.2022.110906

58 Wang, X. et al. Altered mGluR5-Homer scaffolds and corticostriatal connectivity in a Shank3 complete knockout model of autism. Nat Commun 7, 11459 (2016). 10.1038/ncomms11459

59 Bey, A. L. et al. Brain region-specific disruption of Shank3 in mice reveals a dissociation for cortical and striatal circuits in autism-related behaviors. Transl Psychiatry 8, 94 (2018). 10.1038/s41398-018-0142-6

60 Drapeau, E., Riad, M., Kajiwara, Y. & Buxbaum, J. D. Behavioral Phenotyping of an Improved Mouse Model of Phelan-McDermid Syndrome with a Complete Deletion of the Shank3 Gene. eNeuro 5 (2018). 10.1523/eneuro.0046-18.2018

61 Storchi, R. et al. A High-Dimensional Quantification of Mouse Defensive Behaviors Reveals Enhanced Diversity and Stimulus Specificity. Curr Biol 30, 4619–4630.e4615 (2020). 10.1016/j.cub.2020.09.007

62 Blanchard, D. C. & Blanchard, R. J. Ethoexperimental approaches to the biology of emotion. Annu Rev Psychol 39, 43–68 (1988). 10.1146/annurev.ps.39.020188.000355

63 Kopachev, N., Netser, S. & Wagner, S. Sex-dependent features of social behavior differ between distinct laboratory mouse strains and their mixed offspring. iScience 25, 103735 (2022). 10.1016/j.isci.2022.103735

64 Hudry, K. & Slaughter, V. Agent familiarity and emotional context influence the everyday empathic responding of young children with autism. Research in Autism Spectrum Disorders 3, 74–85 (2009).

65 Meyer, M. L. et al. Empathy for the social suffering of friends and strangers recruits distinct patterns of brain activation. Soc Cogn Affect Neurosci 8, 446–454 (2013). 10.1093/scan/nss019

66 Sakurai, K. et al. Capturing and Manipulating Activated Neuronal Ensembles with CANE Delineates a Hypothalamic Social-Fear Circuit. Neuron 92, 739–753 (2016). 10.1016/j.neuron.2016.10.015

67 Davis, M., Walker, D. L., Miles, L. & Grillon, C. Phasic vs sustained fear in rats and humans: role of the extended amygdala in fear vs anxiety. Neuropsychopharmacology 35, 105–135 (2010). 10.1038/npp.2009.109

68 Sah, P. Fear, Anxiety, and the Amygdala. Neuron 96, 1–2 (2017). 10.1016/j.neuron.2017.09.013

69 Gangopadhyay, P., Chawla, M., Dal Monte, O. & Chang, S. W. C. Prefrontal-amygdala circuits in social decision-making. Nat Neurosci 24, 5–18 (2021). 10.1038/s41593-020-00738-9

70 Storchi, R. et al. A High-Dimensional Quantification of Mouse Defensive Behaviors Reveals Enhanced Diversity and Stimulus Specificity. Current Biology 30, 4619–4630.e4615 (2020). 10.1016/j.cub.2020.09.007

71 Wang, S. E. et al. Mechanism of EHMT2-mediated genomic imprinting associated with Prader-Willi syndrome. Res Sq (2024). 10.21203/rs.3.rs-4530649/v1

72 Barczak, W., Suchorska, W., Rubiś, B. & Kulcenty, K. Universal real-time PCR-based assay for lentiviral titration. Mol Biotechnol 57, 195–200 (2015). 10.1007/s12033-014-9815-4

73 Renier, N. et al. iDISCO: A Simple, Rapid Method to Immunolabel Large Tissue Samples for Volume Imaging. Cell 159, 896–910 (2014). 10.1016/j.cell.2014.10.010

